# Macromolecule modelling for improved metabolite quantification using short echo time brain ^1^H MRS at 3 T and 7 T: The PRaMM Model

**DOI:** 10.1101/2023.11.16.567383

**Authors:** Andrea Dell’Orco, Layla Tabea Riemann, Stephen L. R. Ellison, Semiha Aydin, Laura Göschel, Anna Tietze, Michael Scheel, Ariane Fillmer

**Author notes:** Corresponding Author: Andrea Dell’Orco.

## Abstract

**Purpose:** To improve reliability of metabolite quantification at both, 3 T and 7 T, we propose a novel parametrized macromolecules quantification model (PRaMM) for brain ^1^H MRS, in which the ratios of macromolecule peak intensities are used as soft constraints.

**Methods:** Full- and metabolite-nulled spectra were acquired in three different brain regions with different ratios of grey and white matter from six healthy volunteers, at both 3 T and 7 T. Metabolite-nulled spectra were used to identify highly correlated macromolecular signal contributions and estimate the ratios of their intensities. These ratios were then used as soft constraints in the proposed PRaMM model for quantification of full spectra. The PRaMM model was validated by comparison with a single component macromolecule model and a macromolecule subtraction technique. Moreover, the influence of the PRaMM model on the repeatability and reproducibility compared to those other methods was investigated.

**Results:** The developed PRaMM model performed better than the two other approaches in all three investigated brain regions. Several estimates of metabolite concentration and their Cramér-Rao lower bounds were affected by the PRaMM model reproducibility, and repeatability of the achieved concentrations were tested by evaluating the method on a second repeated acquisitions dataset. While the observed effects on both metrics were not significant, the fit quality metrics were improved for the PRaMM method (p≤0.0001). Minimally detectable changes are in the range 0.5 – 1.9 mM and percent coefficients of variations are lower than 10% for almost all the clinically relevant metabolites. Furthermore, potential overparameterization was ruled out.

**Conclusion:** Here, the PRaMM model, a method for an improved quantification of metabolites was developed, and a method to investigate the role of the MM background and its individual components from a clinical perspective is proposed.

## Introduction

MRS is the only non-invasive technique for in-vivo detection and quantification of metabolite concentrations in the human brain. It can provide insight into the neurochemistry of various brain diseases^1^, potentially leading to the discovery of clinically relevant biomarkers^2–6^ and enabling a better understanding of the underlying biochemical processes.

Recently, with the spread of ultra-high-field (≥ 7 T; UHF) MR scanners, it has become possible to acquire spectra with a higher spectral resolution and an increased signal-to-noise ratio^7^ compared to 3 T. Furthermore, the use of an ultra-short echo time (TE) sequence enables the detection and quantification of even more signals^8^, as T2* relaxation and J-coupling effects on the final spectrum are reduced.

However, in ultra-short TE ^1^H-MRS, the complexity of brain spectra is increased by a background of wide peaks stemming from macromolecules (MMs), which were assigned to the resonances of various proteins in previous studies^9–11^. This MM background consists of a series of overlapping, closely spaced, and eventually partially resolved multiplets, originating from the variation of the chemical shift of amino acid protons in different proteins. The same alpha or side chain proton is, in fact, differently shielded from amino acids close in the primary or superior protein structures^12^. This phenomenon results in a continuous distribution of chemical shifts, given the high number of possible combinations^9^. In this way, protons in the same position in an amino acid side chain, but with a different surrounding, may resonate in a chemical shift range of 1 ppm or more. Furthermore, α-protons may resonate in chemical shift ranges of 2 ppm, depending on the protein conformation^12^.

An altered MM background is reported in various diseases^13,14^ and healthy aging^15^, but also depends on the white matter (WM)/gray matter (GM) ratio in the volume of interest (VOI)^16^. However, a conclusive description of the clinical evolution of the biochemistry of brain proteins and their relevance to different diseases is still lacking, as those studies are rare and employ different quantification methods.

Conventionally, the quantification of a ^1^H-MRS spectrum involves a linear combination of a set of basis spectra of individual metabolites or components^17^. To achieve accurate quantification of the metabolites in short TE ^1^H-MRS, the contribution of the MMs must be properly accounted for^18–20^. Approaches to do so, can be roughly divided into two categories: 1) the MM signals are included as a component in the basis set used for linear combination model fitting^21^, which we will refer to here as a single component model (SCM); and 2) the MM signals are subtracted from the full spectrum^22^ usually on an individual basis, which will be referred to as individual metabolite-nulled spectrum subtraction (IMNS) throughout this work.

For SCM, various metabolite-nulled spectra (MNS) are acquired, typically using an inversion recovery or a double inversion recovery (DIR)^9,16^ sequence. The data are then averaged and inserted as a single component in the basis set. A major disadvantage of this method is the lack of flexibility, as it cannot adapt itself to situations where the MM background varies.

To perform IMNS, for each full spectrum (FS), an MNS is measured and subtracted from the FS, resulting in a metabolite-only spectrum^22^. This method suffers from several drawbacks, such as a doubled acquisition time, as for each FS, an MNS must be measured. Moreover, a perfect metabolite nulling is difficult to achieve, leading to a partial subtraction of residual metabolite peaks along with the MMs, which in turn results in biases in the metabolite quantification. Furthermore, subtraction of the MM signals leads to the loss of valuable information about the MMs^9^.

A fitting model, where each MM component is inserted individually in the basis set and quantified together with metabolites, could increase accuracy of metabolite quantification in diseases where the MM background is altered or to investigate MMs as potential disease biomarkers^9^. However, the use of multiple MM components increases the degrees of freedom of the model, and hence, the risk of overparameterization, which would lead to a fit without biochemical meaning^9^.

To avoid overfitting, some prior knowledge or soft constraints are needed^9^. Povazan et al.^10^ suggested the use of concentration ratio priors - expressing the prior probability on chosen intensity ratios^23^ - as soft constraints, using the accurately integrable MM peak at 0.94 ppm as the denominator. Similarly, in the work of Simicic et al. 2021^19^, the soft constraints were based on the signal intensity ratio between each individual Hankel Lanczos Single Value Decomposition (HLSVD) MM component and the easily integrable M0.94 peak^19^.

It is known that intensities of some MM signals are highly correlated with each other because multiple MM peaks can originate either from different protons in the same amino acid or from protons in different amino acids of the same protein, i.e., M0.94 and M3.00 in thymosin-β4^9,24^.

This work aims to develop a parametrized ratio MM (PRaMM) model, where the intensity ratios of highly correlated MM peak intensities are used as soft constraints. Our main hypothesis is that the better modelling of MMs leads to an improved spectral fitting, capable to explain the whole spectral area and, hence, quantification of metabolite concentrations. Moreover, the PRaMM model allows for adaptation to varying MM content, e.g., different GM/WM ratios or potentially diseased tissue. The performance of the proposed model was tested at 3 T and 7 T and compared to the performance of an SCM and IMNS with respect to fit quality number (FQN), residual SNR, and Cramer-Rao Lower Bounds (CRLB). To prevent overfitting, the model was additionally tested for reproducibility and repeatability, and measurement uncertainty and precision of the method were determined.

## Methods

### Instrumentation and participants

Measurements were performed either at a 3 T whole-body MR scanner (MAGNETOM Verio) using a 1Tx/32Rx head coil (both Siemens Healthineers, Erlangen, Germany), or a 7 T whole-body MR scanner (MAGNETOM, Siemens Healthineers, Erlangen, Germany) equipped with a 1TX/32RX head coil (NOVA Medical, Wilmington, USA). Six healthy volunteers were scanned at both field strengths (3 T: age (mean ± SD) 45 ± 14 years; 4:2:0 male:female:non-binary; and 7 T: age (mean ± SD) 50 ± 11 years 5:1:0; female:male:non-binary) for the development of the PRaMM model. At 7 T, 9 additional, healthy volunteers were scanned (age (mean ± SD) 39 ± 13 years; 7:1:1 female:male:non-binary) four times in two sessions, each about one week apart, in order to assess the influence of the PRaMM model on the measurement precision^25^. Hence, in total three data sets were obtained, which will be referred to as the 3 T data set (3T), the 7 T data set (7T), and the repeated acquisition data set (RA) throughout this paper. All volunteers were enrolled after giving written informed consent according to local ethical regulations.

### Double Inversion Recovery sequence

To achieve nulling of metabolites, and hence enable the acquisition of a spectrum containing (ideally) only macromolecular signals, two wideband, uniform rate, smooth truncation (WURST) adiabatic inversion pulses^26^ were added to SPECIAL^27,28^ prior to the localization^28^. The inversion times TI_1_ and TI_2_ were defined as the time interval from the centre of the non-selective inversion pulses to the centre of the excitation pulse. The resulting double-inversion recovery (DIR) sequence will be referred to as DIR-SPECIAL. A schematic representation of the sequence is shown in the supporting material, Fig. S1. The measurement parameters used at 3 T and 7 T are reported in Table 1.

**Table 1:**
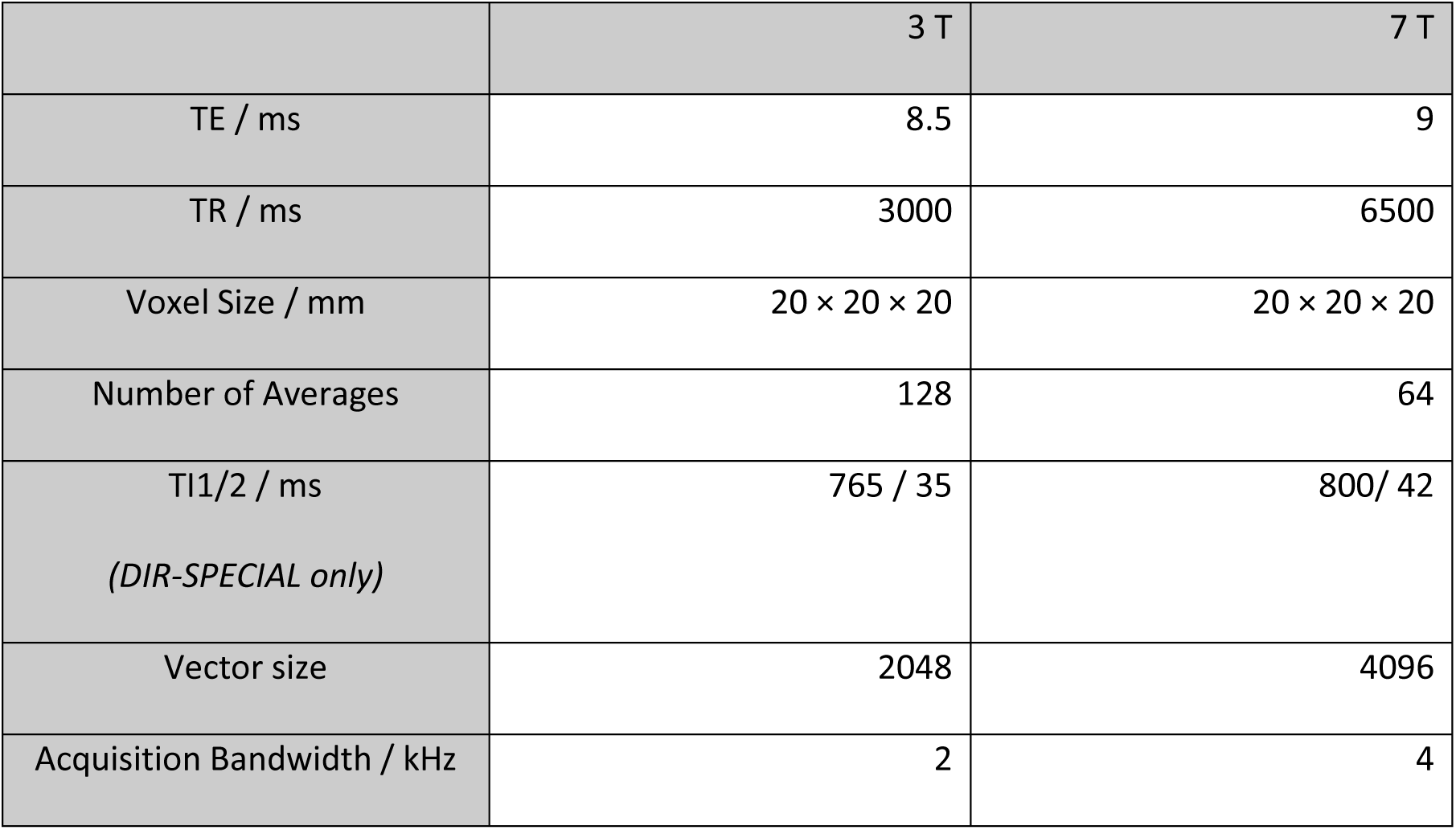
Sequence parameters of the SPECIAL and DIR-SPECIAL sequences, optimized for 3 T and 7 T.

### Phantom Acquisitions

An in-house built spherical phantom containing the following metabolites was used: N-acetyl aspartate (NAA), glutamate (Glu), glutamine (Gln), aspartate (Asp), creatine (Cr), choline (Cho), myo-inositol (mIns). A 0.1 M phosphate buffer at pH 7.2 was used and a 2% agarose gel enriched with monodisperse glass spheres (Monospher 100, Merck, Darmstadt) to simulate microscopic B_0_ inhomogeneities was added. The Bloch equations were employed to simulate the double inversion recovery for 3 T and 7 T, using the formula from Redpath et al.^29^, by fixing a short TI_2_ and calculating an optimal TI_1_. The resulting combination of TI_1_ and TI_2_ was used as starting point to iteratively determine the TI_1_ for metabolite nulling in the phantom. TI_1_ was selected after a visual inspection of the resulting MNS and the intensity of the residual metabolite peaks. As perfect metabolite-nulling was not possible, the residual metabolite peaks were assigned based on the chemical shift and multiplicity of the peaks. The determined combination of TI_1_ and TI_2_ was then used as a starting point for the optimal inversion times in the vivo acquisitions. The effectiveness of the DIR nulling was evaluated by calculating the complement of the ratio of the variance of the MNS to the variance of the full spectrum in the chemical shift range between 4.2 and 0.2 ppm.

### In vivo Acquisitions

#### 3T and 7T

The in vivo acquisitions were similar for both field strengths. T1-weighted MPRAGE^30^ (3T) or MP2RAGE^31^ (7T) were acquired for voxel positioning and subsequent estimation of voxel tissue fractions. After VOI positioning, a localized RF power calibration was performed followed by voxel specific B_0_ shimming with FAST(EST)MAP^32^ and an in-house MATLAB shim tool^33^. After spectral quality assessment by evaluating water linewidth, outer volume saturation bands were positioned adjacent to the voxel, and a full spectrum was acquired using a SPECIAL sequence with the sequence parameters given in Table 1. For absolute quantification a reference measurement was obtained without water suppression with four averages only. Finally, a MNS was acquired using the DIR-SPECIAL sequence. The first in-vivo acquisition at both field strengths was used to find the best TI_1_ and TI_2_ for an optimal nulling of the metabolite signals by varying the inversion times starting with the optimal values obtained in phantom tests. Hence, for those subjects, FS and MNS were acquired only in one Voxel, located in the posterior cingulate cortex (PCC). For all other subjects, FS and MN spectra from two additional voxels were acquired if time permitted, one in the white matter of the parietal lobe alongside the PCC (subsequently abbreviated with WM), and one in the anterior cingulate cortex (ACC). Exemplary VOI positioning is shown in Figure 1.

**Figure 1:**
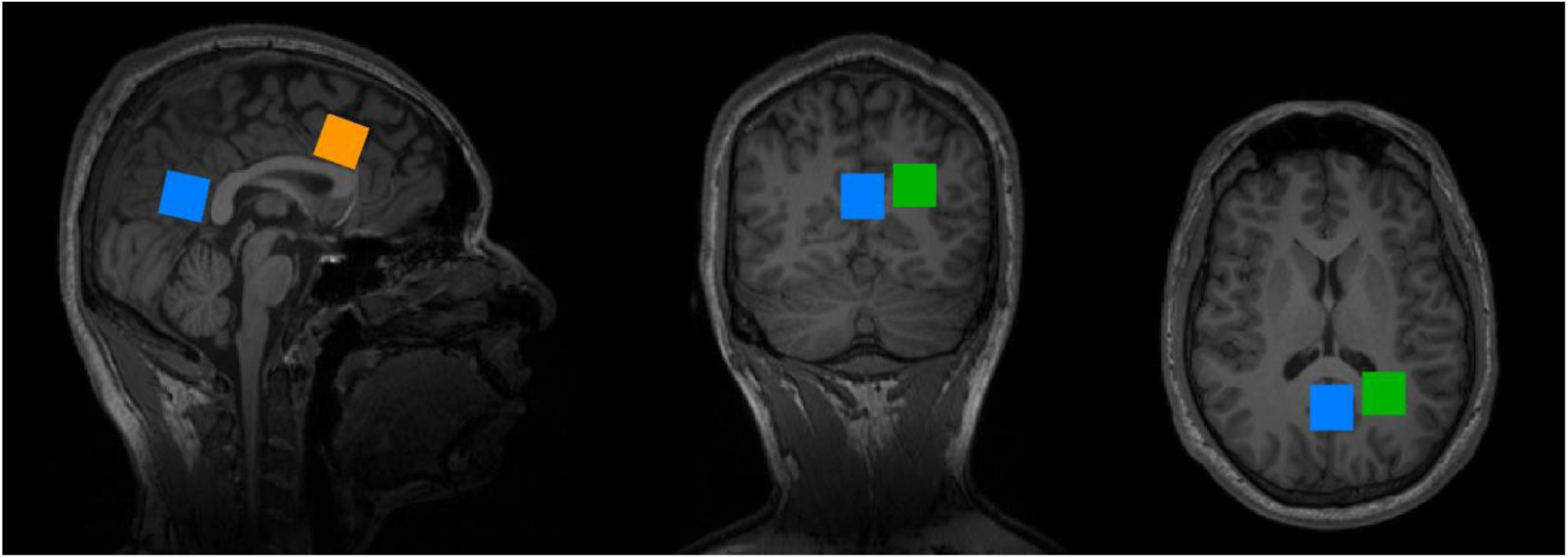
Voxel positioning. Three voxels were acquired: PCC (blue) and ACC (orange), GM-rich areas, adjacent to the corpus callosum, and WM (green) in the WM-rich area of the parietal lobe alongside the PCC.

#### RA

The data for the repeatability data set were obtained according to the unbalanced nested study design described in Rieman et al 2022^25^. Each scan block was composed of the following steps: acquisition of an MP2RAGE image, positioning of the VOI in the PCC, localized RF calibration and B0 optimization, followed by the acquisition of the spectrum and a reference spectrum without water suppression, and using the same SPECIAL sequence as described above.

### Spectral pre-processing

#### 3T and 7T

The spectra were exported from the scanner in Siemens TWIX format and were pre-processed using Python v3.9 with the libraries NumPy v1.18^34^ and Suspect v0.3^35^. The pre-processing pipeline included the following steps: After averaging each odd acquisition with the next, the resulting fully localized FIDs were phase-corrected, and the signals of individual coils combined, employing a weighting based on the single coil SNR^36^. Subsequently, the spectra were frequency corrected and averaged and the residual water signal was removed using the Hankel Lanczos singular value decomposition (HLSVD)^17^. The spectral data were saved in LCModel RAW format.

To obtain the metabolite-only spectra for the IMNS method, further steps were necessary: The negative NAAG peak and, if visible, the Cr methylene at 3.93 ppm were removed in each MNS using a HLSVD with approximately 20 to 40 components. Then, the MNS was scaled to overcome the partial saturation due to the DIR by multiplying it by a factor between 1.2 and 1.5^37^ and finally, each MNS was subtracted from the corresponding full spectrum.

Furthermore, for each voxel position an averaged MNS was calculated, which is used to derive an optimized the MM basis set. The pre-processing code is available on GitHub (https://github.com/0rC0/PRaMM/notebooks).

To determine the tissue composition of each VOI, each spectrum was converted from TWIX format into NIfTI-MRS with spec2nii^38^. A mask of each VOI was generated and applied to the corresponding MP2RAGE images with the svs_segment tool from FSL-MRS^39^.

#### RA

was pre-processed using an in-house developed MATLAB tool including the following steps: summation of odd and even acquisitions, weighted and phase-corrected coil-element combination, frequency correction and averaging. Tissue fractions for the absolute quantification were obtained using an in-house tool for co-registering the voxel position with the MP2RAGE image, which was previously segmented into different tissue types using SPM12^40^.

### Parametrized MM model development

#### 3T and 7T

The averaged MN spectra for both field strengths and the three different brain regions were fitted with LCModel v6.3^41^. For 3 T, the basis set contained only the negative NAAG peak, while at 7 T, the basis set contained also the metabolites revealed in the phantom experiment – NAA and Gln and Glu – and a negative NAAG peak. The MM peaks were simulated as Voigt-lines^42^ using the recommended nomenclature from the expert consensus^9^. Furthermore, at 7T, the Cr methylene and the negative Cr methyl peak at 3.05 ppm were simulated also as Voigt-lines^42^. The MM peaks and their chemical shift are reported in Table 5.

To produce a plausible LCModel fit of the averaged MM spectrum accounting for all MM signals and to avoid the possible attribution of partial MM signals to the baseline^42^, the baseline was forced stiff (DKNTMN = 99). Each MNS was fitted from 4.1 to 0.75 ppm.

After visual inspection of each fit, the parameters relative to the Voigt lines simulating the MM peaks (CHSIMU in LCModel control file), were adjusted and successively optimized to derive the final MM basis set. An example control file can be found on GitHub (https://github.com/0rC0/PRaMM/control_files).

The individual MN spectra were then fitted using the resulting MM basis set. Subsequently, the MM peak intensities were tested for correlations using the Spearman Rank-Order correlation.

Correlations with a rho value higher than 0.66 were considered highly correlated, and the ratios of these correlated peaks were then used as soft constraints in the PRaMM model. Given the sample size of both datasets, the chosen thresholds correspond to a p-value of about 0.01.

Because the prevention of type II errors was considered a priority, a stricter p-value threshold was chosen to limit type I errors, but no p-value adjustment was performed ^41^.

### Spectral quantification

#### 3T, 7T and RA

The spectra were fitted with LCModel^41^ using three different models: 1) the PRaMM model, 2) a SCM model, and 3) an IMNS approach. The basis sets for all three approaches contained the same metabolite basis functions: Alanine (Ala), aspartate (Asp), phosphorylcholine (PCho), creatine (Cr), phosphocreatine (PCr), γ-aminobutyric acid (GABA), glutamine (Gln), glutamate (Glu), glutathione (GSH), glycine (Gly), myo-inositol (Ins), glucose (Glc), lactic acid (Lac), N-acetylaspartate (NAA), scyllo-inositol (Scyllo), taurine (Tau), ascorbic acid (Asc), β-hydroxy-butyrate (BHB), N-acetylaspartylglutamic acid (NAAG), glycerylphosphorylcholin (GPC), phosphorylethanolamine (PE), serine (Ser). Each basis spectrum was simulated with Vespa v0.9.3^43^. The MM signals were treated differently in each model:

1. In the PRaMM model, MM basis functions were included as individual simulated peaks together with the intensity ratios of highly correlated MM peaks from the experiments described above. The information was entered through the LCModel control file using the parameters CHSIMU and CHRATO. A baseline spline node distance of 1 ppm (DKNTMN=1) was chosen as it was found to be optimal in our previous work^44^.
2. For the SCM,a single component MM basis function was used containing all MM peaks in the basis set. The single MM component was obtained by averaging multiple MNS of the PCC, which were acquired using an inversion recovery SPECIAL sequence on multiple healthy volunteers. The DKNTMN was set to 0.25 ppm^19^.
3. For the IMNS approach, pre-processed spectra, from which an individually acquired MN spectrum was subtracted, were fitted with the metabolite-only basis set.

Each spectrum of the 3T and 7T data sets was fitted from 4.1 to 0.2 ppm with each of the methods described above. The RA dataset was only fitted with the SCM and PRaMM model, due to the unavailability of MNS. Water scaling was performed using the LCModel parameters WCONC and ATTH2O and assigning the not water-suppressed reference to the FILH2O parameter.

Because the 3T dataset was incomplete of many MPRAGE images, essential to achieve the water concentration in the VOIs, only the FILH2O parameter was used in the control files.

An example LCModel^41^ control file for each model is available on GitHub (https://github.com/0rC0/PRaMM).

### Absolute quantification of the metabolites

The absolute quantification was performed based on the equations by Near et al. 2020^17^ and as instructed in the LCModel manual, due to an incomplete 3T dataset only for 7T and RA. Briefly, the water signal was considered as the internal standard and its concentration calculated based on the water content and water relaxation times in the individual tissues^17^. The resulting value was assigned to the LCModel WCONC parameter.

The output concentrations from LCModel were further corrected for the CSF fraction and the relaxation times of the individual metabolites. A more detailed explanation is provided in the Supporting Information.

To our best knowledge, no T_2_ values for the MM peaks at 7 T are published yet. For this reason, for the RA at 7T, the MM water-scaled peak intensities were not corrected for relaxation times.

The absolute CRLB (aCRLB) was calculated, multiplying the final concentration or peak intensity value by the relative CRLB (CRLB%).

### Statistical comparisons of the models

#### 3T, 7T, RA

As recommended in a recent experts’ consensus^9^, the fit-quality-number (FQN), was the main metric to compare the quality of the fits. The FQN is defined as the ratio of the variance fit residual and the variance of the pure spectral noise, taken in an area with no signals. Since both the noise-only and the sum of noise and baseline can be considered as the fit residual, we calculate both and refer to them as FQN_n_ and FQN_bln_, respectively.

The same value of pure spectral noise, taken at negative chemical shifts, was used for all models. The SNR that was extracted from the LCModel COORD file, was used as a further metric. It is defined in LCModel as the ratio of the maximum in the spectrum-minus-baseline over the fitting range to twice the residual^45^.

The effect of the fitting model on the concentrations of the main metabolites (7T) or water-scaled peak intensities (3T), and the absolute Cramèr-Rao lower bounds (aCRLB) were statistically tested as follows: As it was not possible to assume a normal distribution of the measurements, non-parametric tests were used. Friedman test for repeated acquisitions was used for group comparison and a paired Wilcoxon signed-rank test^46^ as a post hoc test for 3T and 7T; statistical tests on RA were done using the paired Wilcoxon signed-rank test. In the Friedman group comparison, the effect size, expressed as Kendall W^47^, was calculated as well. The effect size must be considered large when higher or equal to 0.5, moderate between 0.3 and 0.5, and small when lower than 0.3. To avoid confusion between the Kendall W and the statistical W value from the Wilcoxon signed-rank post hoc test, the two parameters are indicated as W_K_ and W_W_, respectively. Where the Friedman’s test condition of “unreplicated complete block design” was not satisfied, only the paired Wilcoxon signed-rank test was applied.

The effect of the method on the repeatability and reproducibility of the derived metabolite concentration was estimated by calculating the variance component for each metabolite and method using a restricted maximum likelihood estimation (REML)^48^. The significance of the post-processing effects on concentrations was checked via post hoc pairwise comparison – using Tukey’s test - applied to the mixed-effect model. Furthermore, the minimal detectable change (MDC) was calculated using the standard error of measurement (SEM)^25^ and the coefficient Variation (CV) was calculated as well as the ratio of standard deviation and mean within all sessions of each subject.^20^ We will consider a percent coefficient variation (CV_%_) ≤ 10% as very good, a CV_%_ ≤ 20% as acceptable, CV_%_ > 20% as not acceptable.

A p-value ≤ 0.05 was considered significant. The Bonferroni adjustment of the p-value was used as multiple-test correction for the main test, Bonferroni-Holm for post hoc. Adjusted p-values are reported as p_adj_.

## Results

The final data set contains 6 PCC acquisitions, 5 WM acquisitions, and 3 ACC acquisitions, resulting in a total number of 14 FS and 13 MNS for 3T, as well as 5 PCC acquisitions, 5 WM acquisitions, and 4 ACC acquisitions, leading to a total of 14 FS and 13 MNS for 7T. One spectrum was excluded from the RA dataset due to poor quality in one measurement, leading to a total of 32 PCC spectra from 8 subjects.

### DIR Sequence and phantom tests

The Bloch equation simulation predicted TI_1_ = 650 ms and TI_1_ = 765 ms for the 3 T and 7 T, respectively, while TI_2_ was set to 35 ms in both cases. Through successive optimizations, the best combination of TI_1_ and TI_2_ for the phantom measurement was determined to be 750 ms and 30 ms for 3 T, and 800 ms and 45 ms for 7 T. The effectiveness of the DIR nulling was 99.40% and 97.92% for 3 T and 7 T, respectively.

At 3 T, the spectrum of the phantom showed residual peaks assignable to multiple metabolites and several small, negative peaks in the 2.5 – 2.2 ppm range, which were tentatively assigned to the resonance of the methylene groups of Glu and Gln, as well as the residual peak of the Cr methyl group at 3.00 ppm.

At 7 T, the residual of the whole spectra of NAA, Glu, and Gln as well as the negative peak of the trimethylamino group of Cho were identifiable.

A comparison between full- and metabolite-nulled spectra for phantom acquisitions at 3 T and 7 T is shown in Supporting Information Figure S2.

### In-vivo measurements

Figure 2 shows spectra acquired with SPECIAL (Fig. A, C) and DIR-SPECIAL (Fig. B, D) sequence at different inversion times at 3 T (A, B) and 7 T (C, D). A high spectral quality was achieved with an average water linewidth of 6.54 ± 0.45 Hz and 12.57 ± 1.34 Hz at 3 T and 7 T, respectively. TI_1_ and TI_2_ were chosen to be set at 765 ms and 35 ms at 3 T, and 800 ms and 42 ms at 7 T for the subsequent in vivo measurements. Average tissue fractions for the different VOI positions are reported in Supporting Information Table S1.

**Figure 2:**
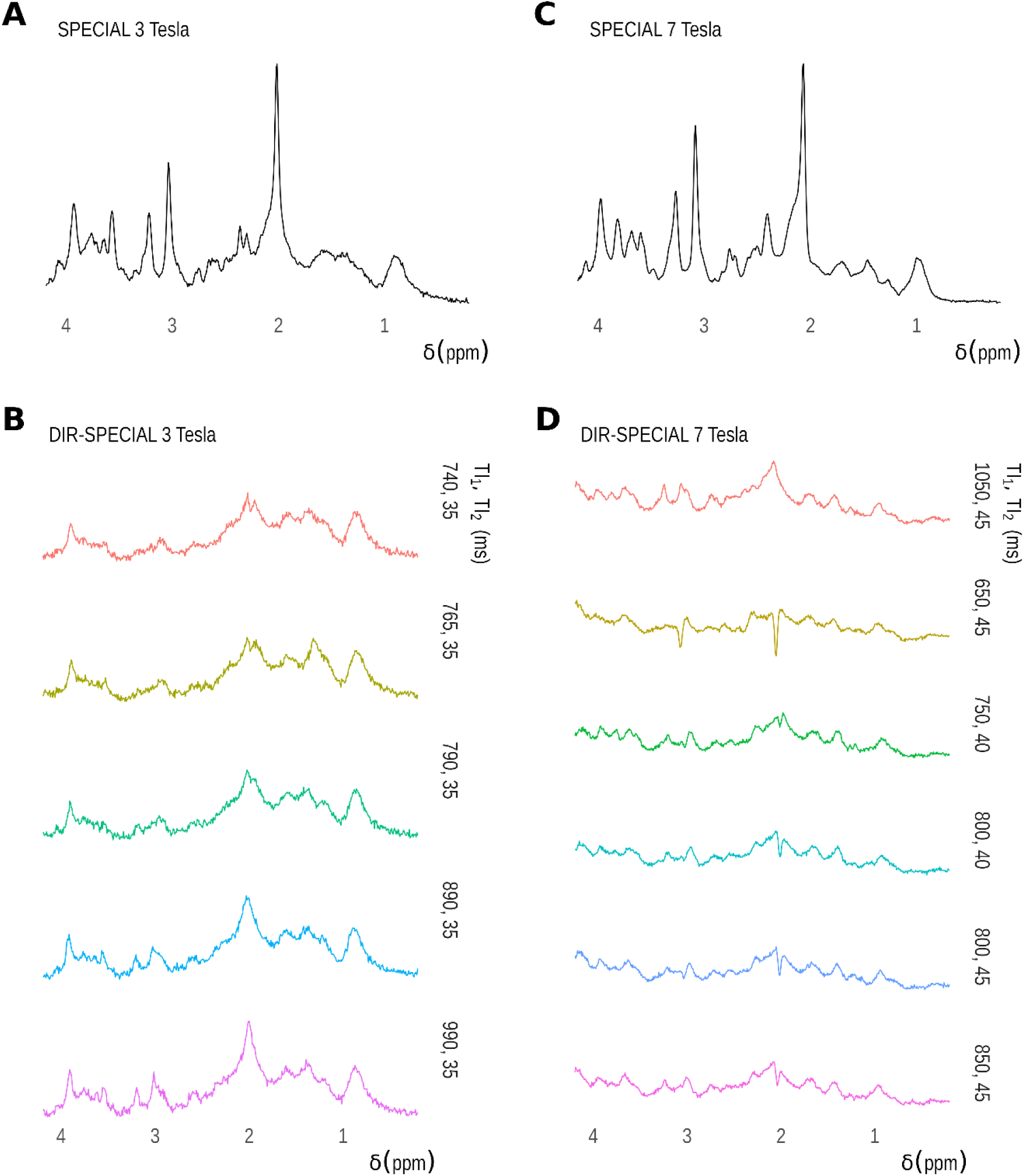
In-vivo spectra from SPECIAL (A, C) and DIR-SPECIAL (B, D) acquired at 3 Tesla (A, B) and 7 Tesla (C, D). The selected DIR-SPECIAL inversion time combination was TI1/TI2 of 765/35 ms at 3T and 800/42 ms at 7T. This choice represents a compromise between residual positive peaks (e.g., Cr at 3.00 ppm) and negative peaks (e.g., NAAG at 2.10 ppm). In the 7T series, the NAAG peak appears negative for almost all combinations.

### Macromolecular basis set and the PRaMM model

The characterization of the MNS led to the identification of the 12 MM peaks for 3T and 15 MM peaks for 7T. The individual MM peaks along with the averaged MNS for both field strengths are displayed in Figure 3 and their intensities are reported in Table 2^34^ Figure 3 shows an exemplary individual MNS for the ACC (a, b), PCC (c, d), and WM (e, f), for 3 T (a, c, e) and 7 T (b, d, f), along with the LCModel fits using the derived macromolecular basis set.

**Figure 3:**
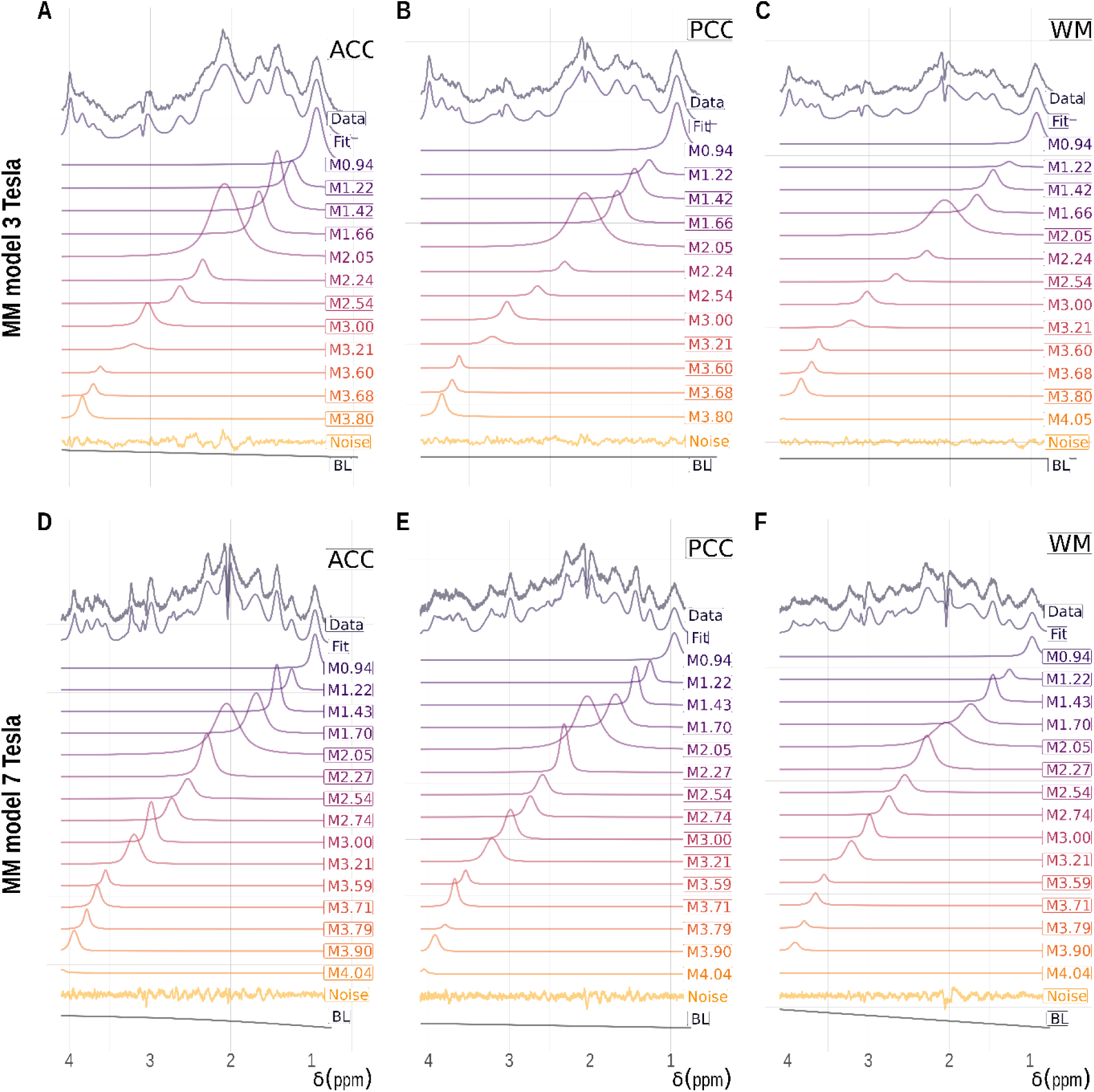
Exemplary individual MNS and simulated fitted MM peaks for the ACC (A,D), PCC (B,E), WM (C,F) at 3 T (A,B,C) and 7 T (D,E,F). At both fields the negative NAAG and Cr peaks are visible in almost all VOIs. The Cr methylene at 3.92 is more intense at 3 T and efficiently nulled at 7 T. BL: Baseline

**Table 2:**
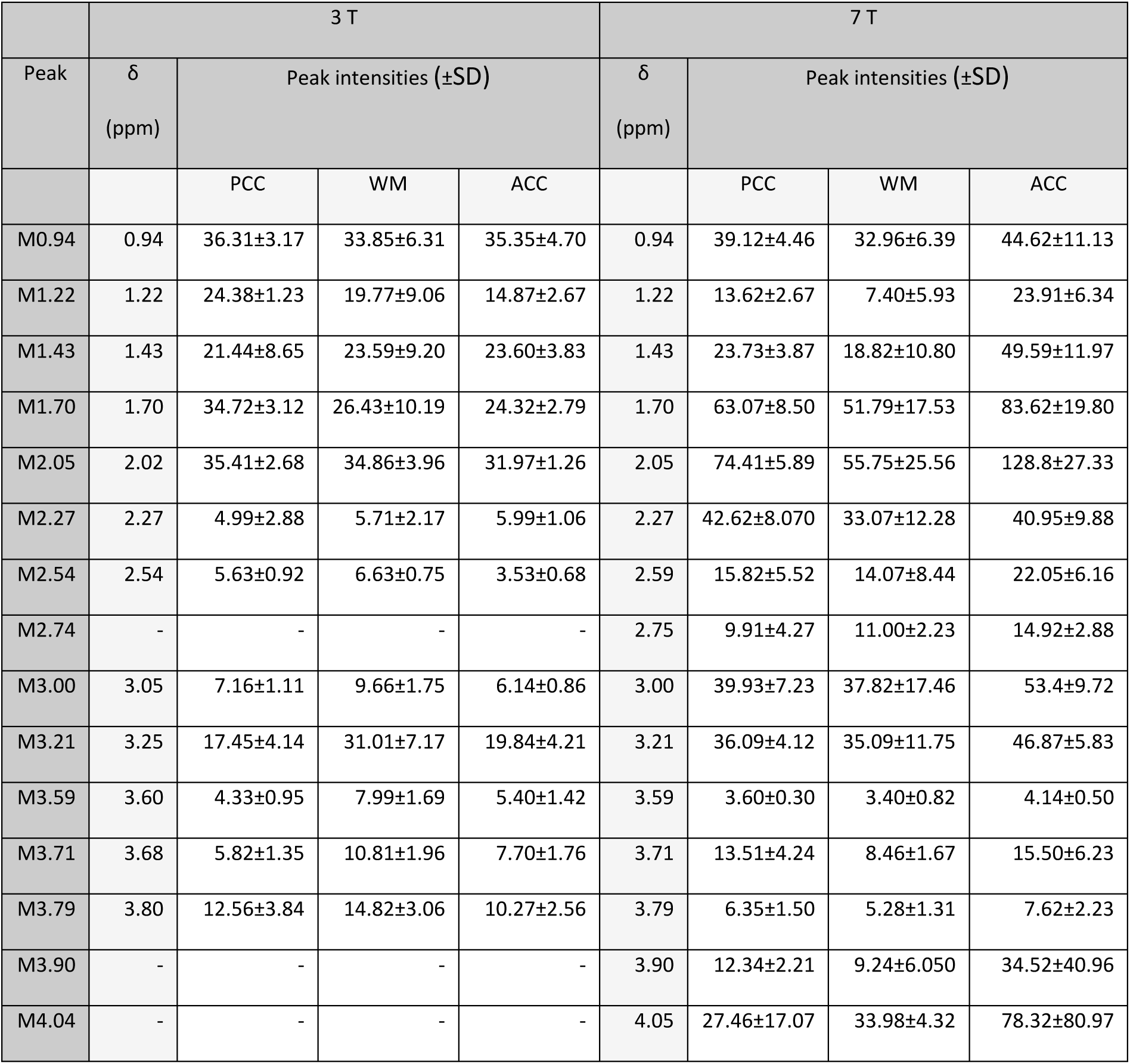
Identified MM peaks, their chemical shifts and mean peak intensities (±SD) for 3 T and 7 T for all investigated brain regions (PCC, ACC, WM) in the arbitrary units from LCModel. The partial volume correction was performed only at 7 T due to the lack of anatomical data at 3 T. The missing values at 3 T represent the peaks found at 7 T but not at 3 T.

After individual fitting of all MNS spectra, peak intensities were tested for Spearman Rank-Order correlation and 12 and 14 couples of highly correlated MM peaks were found in the 3T and 7T spectra, respectively. The intensity ratios of highly correlated MM peaks, which were subsequently used as soft constraints in the PRaMM model, and the relative Spearman’s ρ are reported in Table 3.

**Table 3:**
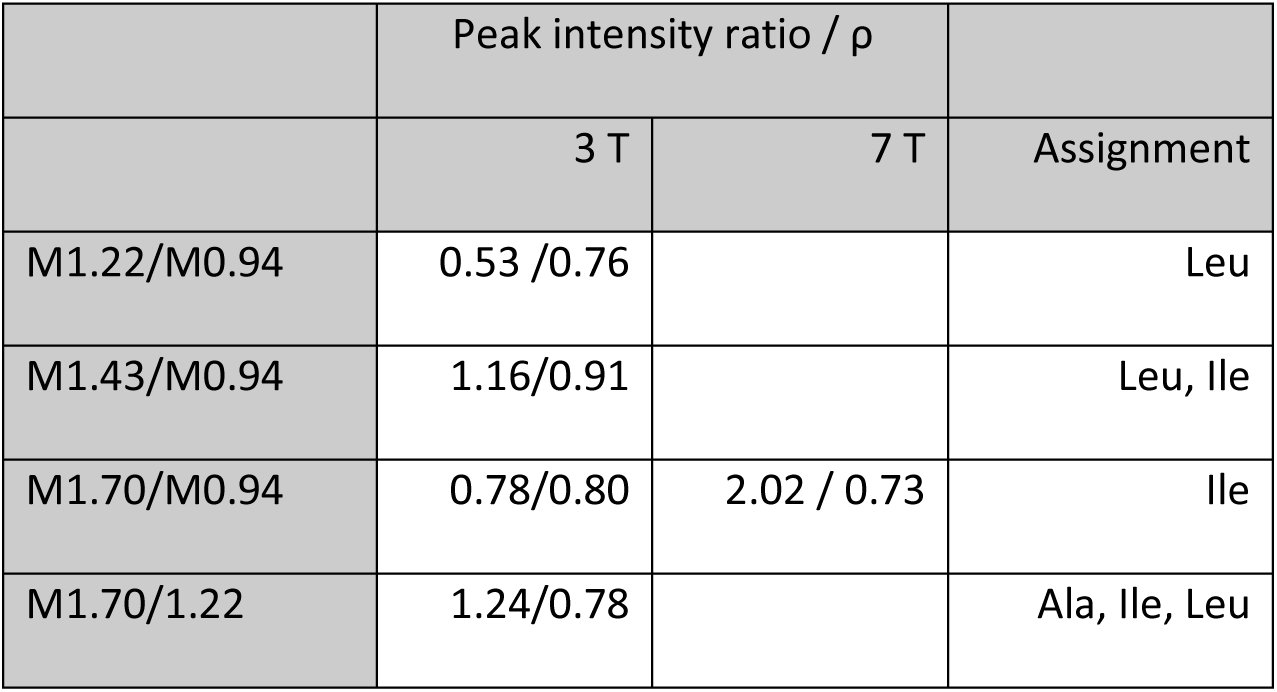
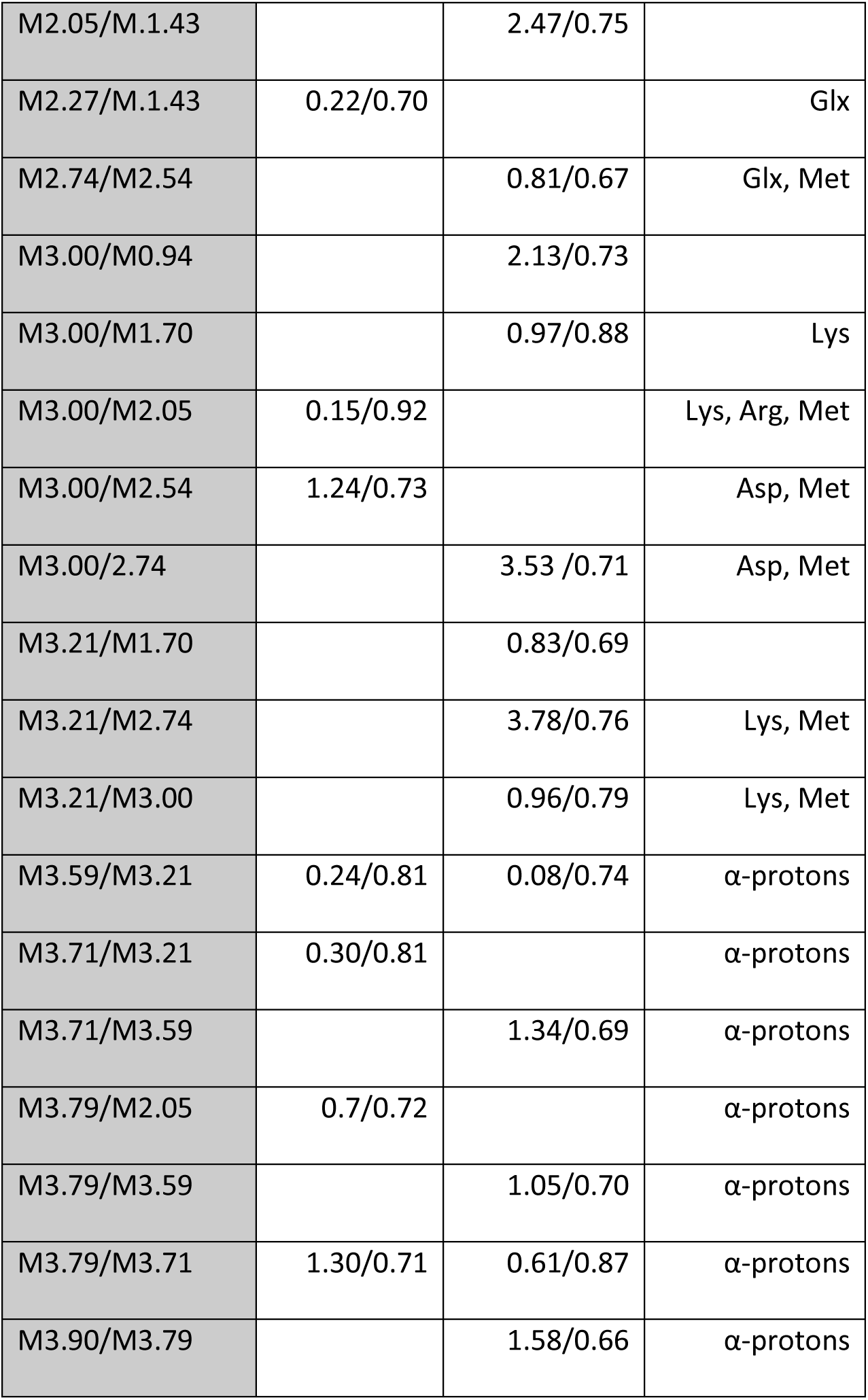
Couples of highly correlated MM peaks and corresponding tentative assignments for 3T and 7T based on Biological Magnetic Resonance Bank (BMRB)^50^.

| | Peak intensity ratio / $\rho$ | | Assignment |
| --- | --- | --- | --- |
|  | 3 T | 7 T |  |
| M1.22/M0.94 | 0.53 / 0.76 |  | Leu |
| M1.43/M0.94 | 1.16/0.91 |  | Leu, Ile |
| M1.70/M0.94 | 0.78/0.80 | 2.02 / 0.73 | Ile |
| M1.70/1.22 | 1.24/0.78 |  | Ala, Ile, Leu |
| M2.05/M.1.43 |  | 2.47/0.75 |  |
| M2.27/M.1.43 | 0.22/0.70 |  | Glx |
| M2.74/M2.54 |  | 0.81/0.67 | Glx, Met |
| M3.00/M0.94 |  | 2.13/0.73 |  |
| M3.00/M1.70 |  | 0.97/0.88 | Lys |
| M3.00/M2.05 | 0.15/0.92 |  | Lys, Arg, Met |
| M3.00/M2.54 | 1.24/0.73 |  | Asp, Met |
| M3.00/2.74 |  | 3.53 /0.71 | Asp, Met |
| M3.21/M1.70 |  | 0.83/0.69 |  |
| M3.21/M2.74 |  | 3.78/0.76 | Lys, Met |
| M3.21/M3.00 |  | 0.96/0.79 | Lys, Met |
| M3.59/M3.21 | 0.24/0.81 | 0.08/0.74 | α-protons |
| M3.71/M3.21 | 0.30/0.81 | | α-protons |
| M3.71/M3.59 | | 1.34/0.69 | α-protons |
| M3.79/M2.05 | 0.7/0.72 | | α-protons |
| M3.79/M3.59 | | 1.05/0.70 | α-protons |
| M3.79/M3.71 | 1.30/0.71 | 0.61/0.87 | α-protons |
| M3.90/M3.79 | | 1.58/0.66 | α-protons |

### Influence of MM handling on Spectral Quantification

Averaged noise and baseline from LCModel quantification for each model and VOI and for both field strengths are shown in Figure 4. Results from statistical tests on concentrations are presented in Table 4. Peak intensities for 3T and metabolite concentrations for 7T are stated in Table 5.

**Figure 4:**
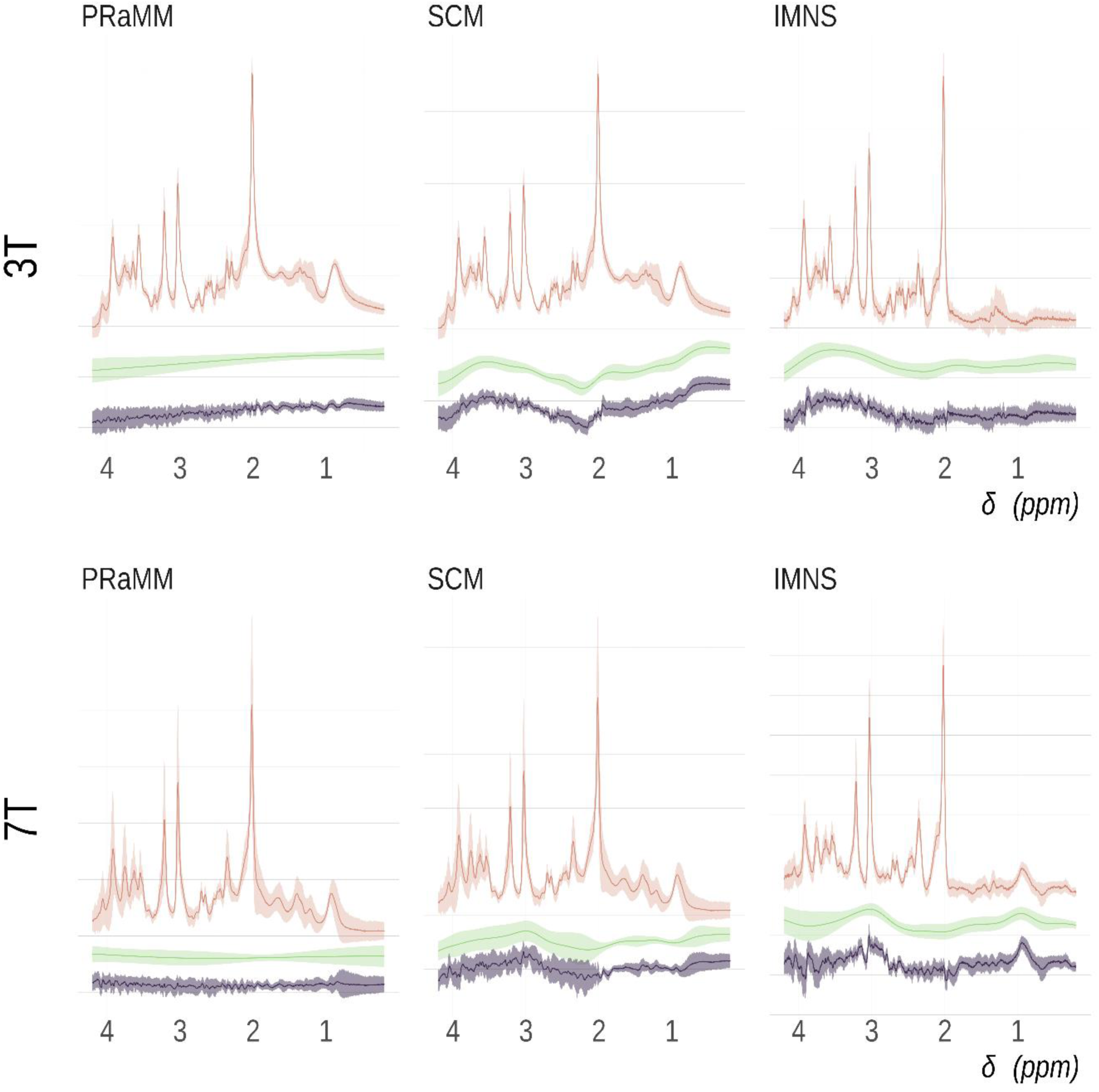
Averaged spectra (orange), baseline (green) and fit residual (violet) averaged across all subjects and VOI localizations for 3 T (above) and 7 T (below) for all investigated MM treatments. Shaded areas represent the standard deviations. Although forcing the baseline flat, as done when using the PRaMM model, may increase the residual, with the PRaMM model a lower residual was achieved than with the other models. Using IMNS, the fits exhibit better baseline at 3 T than at 7 T, where two local maxima can be seen due to the suboptimal nulling of residual metabolite signals and partial saturation of M0.95 and M3.00. This demonstrates that at 7 T it could be even more challenging to obtain an optimal MNS. In the SCM fits, at both 3 T and 7 T, multiple local maxima and minima can be observed in the baseline in correspondence with the chemical shifts of major MM peaks, indicating a suboptimal assignment of MM signals.

**Table 4:** results of statistical testing on peak intensities, concentrations, and aCRLB for all three datasets. The Friedman test for CRLB was not performed due to the unmet requirement of an “unreplicated block design”. The initial test underwent Bonferroni correction, followed by Bonferroni-Holm correction in post-hoc analysis. Abbreviations: -: not tested; ns: p_adj_ >0.05, *: p_adj_ ≤0.05, **: p_adj_ ≤0.01, ***: p_adj_ ≤0.001, ****: p_adj_ ≤0.0001.

| Metabolite | Friedman's test ($\chi^2$, $p_{adj}$, $W_k$) | | | | | | |
| --- | --- | --- | --- | --- | --- | --- | --- |
|  | 3T |  |  | 7T |  |  |  |
| GABA | - |  |  | 3,5 | ns | 0,145833 |  |
| Gln | 16,77 | ** | 0,64 | 4,666667 | ns | 0,194444 |  |
| Glu | 12,92 | * | 0,50 | 12,5 | * | 0,520833 |  |
| GSH | 9,38 | ns | 0,36 | 15,16667 | ** | 0,631944 |  |

| Ins | 7,54 | ns | 0,29 | 8,166667 | ns | 0,340278 |  |
| --- | --- | --- | --- | --- | --- | --- | --- |
| Lac | 11,02 | * | 0,42 | 5,166667 | ns | 0,215278 |  |
| tCho | 19,85 | *** | 0,76 | 8,666667 | ns | 0,361111 |  |
| tCr | 26,00 | **** | 1,00 | 11,16667 | * | 0,465278 |  |
| tNAA | 19,85 | *** | 0,79 | 4,166667 | ns | 0,173611 |  |
| Pairwise Wilcoxon signed-rank test, post-hoc ($W_w$, $p_{adj}$) | | | | | | | |
| Metabolite | 3T |  |  | 7T |  |  | RA |
|  | PRaMM vs SCM | PRaMM vs IMNS | SCM vs IMNS | PRaMM vs SCM | PRaMM vs IMNS | SCM vs IMNS | PRaMM vs SCM |
| GABA | 91 ** | 91 ** | 61 ns | - | - | - | 0 ** |
| Gln | 87 * | 31 ns | 0 ** | - | - | - | 45 ** |
| Glu | 11 ns | 4 ns | 19 ns | 48 ns | 1 * | 2 * | 22 ns |
| GSH | - | - | - | 0 ** | 4 ns | 40 ns | 23 ns |
| Ins | - | - | - | - | - | - | 45 ** |
| Lac | 2 ns | 1 ns | 6 ns | - | - | - | 45 ** |
| tCho | 0 ** | 0 ** | 52 ns | - | - | - | 9 ns |
| tCr | 0 ** | 0 ** | 0 ** | 7 ns | 3 ns | 11 ns | 0 ** |
| tNAA | 0 ** | 0 ** | 67 ns | - | - | - | 0 ** |
| Pairwise Wilcoxon signed-rank test($W_w$, $p_{adj}$) | | | | | | | |
| aCRLB | 3T |  |  | 7T |  |  |  |
|  | PRaMM vs SCM | PRaMM vs IMNS | SCM vs IMNS | PRaMM vs SCM | PRaMM vs IMNS | SCM vs IMNS |  |
| GABA | 0 ** | 0 ** | 35 ns | 45 * | 28 ns | 0 ** |  |
| Gln | 1 ** | 23 ns | 83 * | 5 ns | 5 ns | 3 ns |  |
| Glu | 33 ns | 47 ns | 22 ns | 45 * | 22 ns | 0 * |  |
| GSH | 62 ns | 17 ns | 0 ** | 78 ** | 75 ** | 0 * |  |
| Ins | 55 ns | 22 ns | 10 ns | 0 ns | 0 ** | 0 ** |  |
| Lac | 53 * | 55 * | 40 ns | 12 ns | 28 ns | 47 ns |  |
| tCho | 60 ns | 166 ns | 100 ns | 66 * | 39 * | 28 ns |  |
| tCr | 171 *** | 253 ***** | 37 ns | 28 * | 28 ns | 14 ns |  |
| tNAA | 36 * | 41 ns | 1.5 ns | 27 ns | 2 ns | 0 ns |  |

**Table 5:** Metabolite concentration results from PRaMM LCModel quantification and comparison of results with VOIs in similar positions from previous studies.

| Metabolite | Absolute concentrations / mM |  |  |  |  |  |  |
| --- | --- | --- | --- | --- | --- | --- | --- |
|  | This paper, PRaMM 7T, |  |  | Marjanmska et al. 2019 |  | Giapitzakis et al 2018 |  |
|  | ACC | PCC | WM | PCC | OCC | Occipital | Left Parietal |
| Asp | 7.8±2.12 | 3.84±0.85 | 2.49±1.92 | 1.98±0.06 | 1.75±0.06 | 2.3±0.5 | 1.5±0.8 |
| GABA | 4.92±0.74 | 2.74±0.41 | 2.08±0.74 | 0.87±0.04 | 1.04±0.04 | 0.8±0.2 | 0.8±0.4 |
| Gln | 9.15±1.75 | 4.81±0.3 | 3.51±1.37 | 2.40±0.98 | 3.35±0.07 | 2.97±0.07 | 1.7±0.4 |
| Glu | 22.1±2.82 | 13.86±1.79 | 9.71±1.12 | 8.80±0.75 | 10.13±0.16 | 8.95±0.16 | 5.8±0.4 |
| GSH | 0.98±1.11 | 1.18±1.17 | 0.9±0.78 | 1.01±0.22 | 1.14±0.02 | 1.05±0.02 | 1.5±0.2 |
| Ins | 15.54±3.81 | 8.32±1.85 | 8.29±1.01 | 7.69±0.18 | 7.69±0.18 | 7.06±0.18 | 4.7±0.9 |
| Lac | 2±1.58 | 1.08±0.46 | 0.48±0.19 | - | - |  | 0.4±0.3 |
| NAA | 19.36±1.12 | 13.57±2.01 | 11±0.59 | 10.80±0.88 | 10.84±0.19 | 11.65±0.19 | 7.9±0.7 |
| NAAG | 3.58±0.49 | 2.25±0.17 | 4.17±0.75 | 4.20±0.92 | 0.99±0.03 | 1.07±0.03 | 1.1±0.3 |
| PE | 4.47±2.09 | 3.22±0.64 | 1.67±1.33 | 1.05±0.90 | 1.16±0.04 | 0.77±0.04 | 0.7±0.3 |
| Tau | 0.51±0.85 | 0.72±0.62 | 0.11±0.21 | 0.89±0.63 | 1.83±0.05 | 1.80±0.03 | 1.1±0.7 |
| tCho | 3.56±0.94 | 0.95±0.4 | 2.16±0.54 | 1.47±0.03 | 1.16±0.03 |  | 1.3±0.2 |
| tCr | 15.56±2.19 | 9.13±2.09 | 7.59±1.19 | 9.70±0.17 | 9.70±0.17 |  | 6.4±0.6 |

#### 3T

The Friedman test revealed significant differences in metabolite peak intensity estimates among the three models, specifically for GABA, Gln, Glu, Lac, tCho, tCr, and tNAA.

Post hoc comparisons indicated significant differences between PRaMM and SCM for GABA, Gln, tCho, tCr, and tNAA. Differences between PRaMM and IMNS were found for GABA, tCho, tCr, and tNAA. SCM exhibited significant differences with IMNS only for Gln and tCr.

Pairwise Wilcoxon signed-rank tests on the aCRLB revealed significant differences between PRaMM and SCM for GABA, Gln, Lac, tCr, and tNAA; between PRaMM and IMNS for GABA, Lac, and tCr; and between SCM and IMNS for Gln and GSH., between PRaMM and IMNS for GABA, Lac and tCr, between SCM and IMNS for Gln and GSH.

#### 7T

The Kruskal-Wallis test showed no significant effect of the VOI on the MM estimates. However, effect sizes were large for M0.94, M1.22, M1.43, M1.70, M2.05, M2.74, M3.21, and M3.71; moderate for M2.54, M3.00, M3.59, and M3.90; and small for M2.27, M3.79, and M4.04.

The Friedman test showed significant differences, post-Bonferroni adjustment, in the absolute concentrations of Glu, GSH, and tCr. Post hoc comparisons revealed significant differences between PRaMM and SCM for GSH and between PRaMM and IMNS, as well as SCM and IMNS, for Glu.

Pairwise Wilcoxon signed-rank tests on the aCRLB showed significant differences between PRaMM and SCM for GABA, Glu, GSH, tCho, and tCr; between PRaMM and IMNS for GSH, Ins, and tCho; and between SCM and IMNS for GABA, Glu, GSH, and Ins.

#### RA

The paired Wilcoxon test indicated a significant difference between PRaMM and SCM for GABA, Gln, Ins, Lac, tCr, and tNAA.

### Statistical comparisons of the models

#### 3T

Friedman test showed significant FQN differences between the models for both FQN_n_ (χ^2^ = 19.53, p_adj_ ≤0.0001, W_K_=0.75, Figure 5A) and FQN_bln_ (χ^2^ =18, p_adj_≤0.001, W_K_=0.69, Figure 5B). For the first case, the pairwise Wilcoxon signed rank test revealed a significantly lower FQN_n_ for PRaMM than for IMNS (W_W_=0, p_adj_≤0.0001), while SCM performed better than IMNS (W_W_=0, p_adj_≤0.0001). In the latter case, the difference in FQN_bln_ was significant between PRaMM and SCM (W_W_=0, p_adj_≤0.01) and between SCM and IMNS (W_W_=89, p_adj_≤0.001). When testing the SNR (χ^2^ =23.8, p_adj_≤0.0001, Figure 5C), SCM showed a significantly higher SNR than PRaMM (W_W_=0, p_adj_≤0.01) and IMNS (W_W_=91, p_adj_≤0.01). PRaMM showed a significantly higher SNR than IMNS (W_W_=76, p_adj_≤0.05).

**Figure 5:**
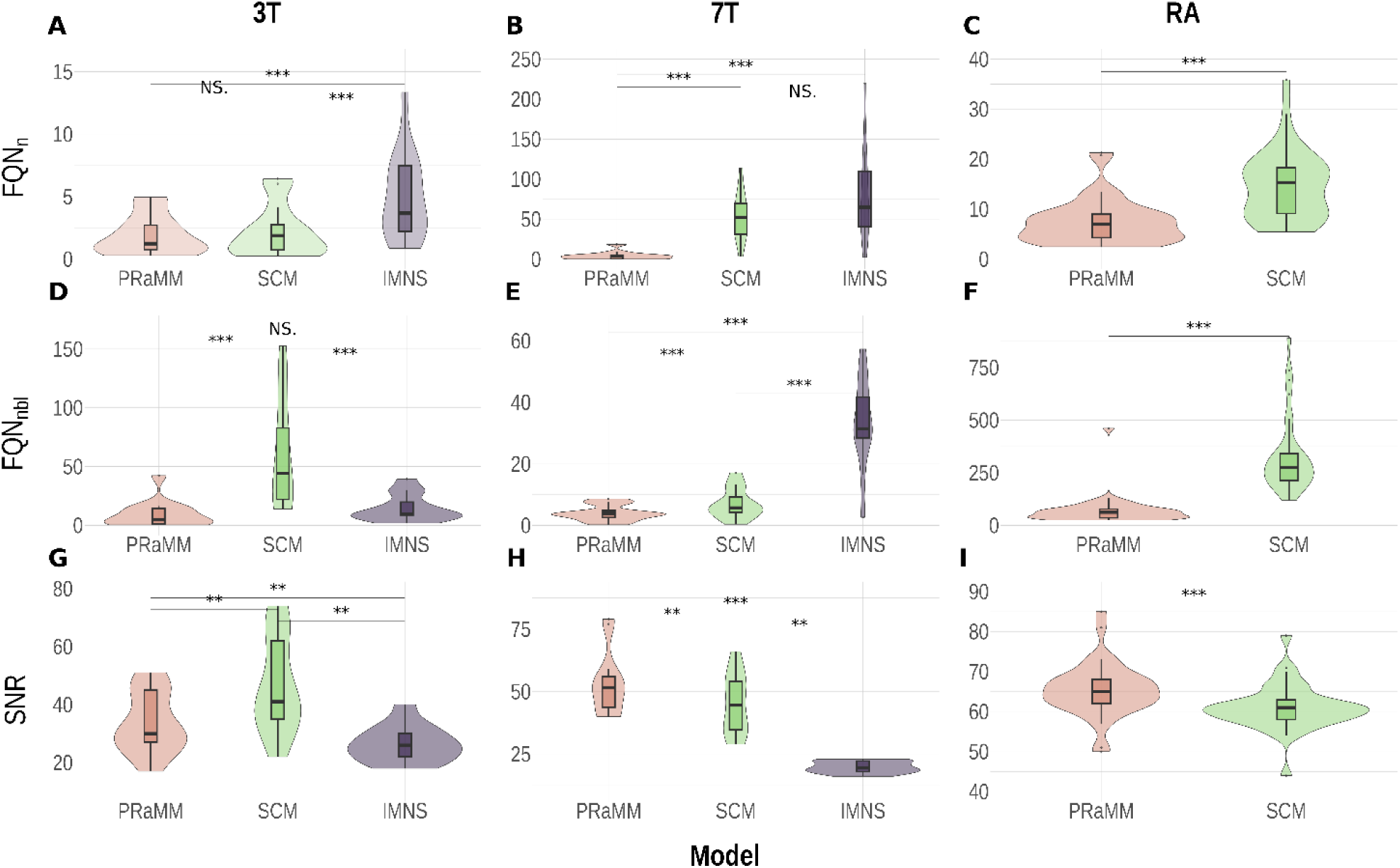
Boxplots and value distributions for the fit quality parameters FQN_n_ (A, D, G), FQN_bln_ (B, E, H), and SNR (C, F, I) for all three acquisitions. With respect to the FQN, the PRaMM model exhibits an improved performance compared to the other two models, although not significant at 3T. With respect to SNR, it appears that the SCM leads to better results, than the PRaMM model. However, it should be noted, that the SNR used for this comparison is extracted from LCModel and biased by the local minimum in the LCModel baseline located below the NAA peak. Notes: G,H,I the y axes do not start from 0.

#### 7T

Friedman test was significant both in the case FQN_n_ (χ^2^ =24.00, p≤0.0001, W_k_=1, Figure 5D) and in the case of FQN_bln_ (χ^2^ =18.66, p≤0.0001, W_k_=0.77, Figure 5E). Post hoc pairwise test revealed significantly improved performances of the PRaMM model with respect to SCM (W_W_=0, p_adj_≤0.001) and IMNS (W_W_=0, p_adj_≤0.001). Additionally, SCM outperformed IMNS (W_W_=0, p_adj_≤0.001) in the first case. In the latter scenario, significant differences were observed between PRaMM and SCM (W_W_=0, p_adj_≤0.001), and between PRaMM and IMNS (W_W_=0 p_adj_≤0.001).

The Friedman test on the SNR showed an effect of the model (χ^2^ =21.82, p_adj_≤0001, W_K_=0.90, Figure 5F): The PRaMM model showed a higher (better) SNR than SCM (W_K_=65, p_adj_≤0.05) and IMNS (W_K_=78, p_adj_≤0.01). Additionally, SCM outperformed IMNS (W_K_=78, p_adj_≤0.01). Boxplots representing these metrics are shown in Figure 5.

#### RA

Paired Wilcoxon test revealed better performances of PRaMM with FQNn (W_W_=0, p_adj_≤0.0001), FQNbln (W_W_=34, p_adj_≤0.0001), and SNR (W_W_=557, p_adj_≤0.0001).

The Friedman test on the REML analysis showed no significant effect of the quantification method neither on reproducibility nor on repeatability. SEM and MDC are reported in Table 6, the SD components for each metabolite and their CVs in Figure 8.

**Table 6:** Minimally Detectable Change and percent coefficient variation (CV_%_) for both PRaMM and SCM. No consistent trend can be observed, which of the models leads to a better performance. However, it should be noted that SCM uses a MM background measured in the same Voxel position in healthy volunteers, as the RA dataset, which might positively bias the performance of SCM in this comparison.

| Metabolite | MDC -<br>PRaMM<br>(mM) | MDC -<br>SCM<br>(mM) | CV% - PRaMM | CV% - SCM |
| --- | --- | --- | --- | --- |
| Asp | 1.529 | 1.196 | 15.51±7.43 | 7.59±3.79 |
| GABA | 0.850 | 1.247 | 11.24±3.71 | 11.08±5.18 |
| Gln | 0.918 | 0.641 | 5.89±1.89 | 4.42±2.38 |
| Glu | 1.605 | 2.012 | 3.81±1.30 | 4.52±2.11 |
| GSH | 0.592 | 0.477 | 10.97±6.66 | 9.56±4.17 |
| Ins | 1.944 | 1.064 | 6.12±3.68 | 4.39±4.02 |
| Lac | 0.800 | 0.375 | 22.34±10.38 | 48.67±10.38 |
| NAA | 1.503 | 1.794 | 3.30±1.84 | 3.62±2.3 |
| NAAG | 0.404 | 0.612 | 5.84±3.01 | 11.21±6.27 |
| Tau | 0.702 | 0.770 | 43.25±45.02 | 31.52±23.22 |
| tCho | 0.555 | 0.234 | 11.50±6.22 | 4.99±2.21 |
| tCr | 1.155 | 1.023 | 3.40±3.27 | 3.37±1.38 |

## Discussion

In this work, we propose the PRaMM model for improved quantification of ultra-short TE proton MRS spectra. The model was applied to the quantification spectra acquired at 3 T and 7 T and led to an improved fit compared to two other conventional fitting models (SCM and IMNS) in most metrics. The PRaMM model was validated on a new 7 T dataset and the improved fit was confirmed. The effect of the model on the reproducibility and repeatability of the estimates of metabolite concentrations exhibited no significant differences in comparison with SCM.

The implemented DIR-SPECIAL sequence with adjusted inversion times reliably nulled most of the metabolite signals at both 3 T and 7 T. While overall a high quality of MNS was achieved, a negative peak appeared at approximately 0.5 ppm in the spectra acquired at 7 T. This is most likely an artifact and can be attributed to partial saturation effects. To avoid any detrimental influence of this on the developed model, the fitting range was restricted to 0.75 - 4.1 ppm in LCModel. Nevertheless, the fit of M0.94 in the full spectra applying the proposed PRaMM model resulted in low relative CRLB. On the other hand, using the IMNS method, the saturation effects led to a notably influenced fit residual, as it was not possible to subtract the M0.94 completely. This explains the poorer performance of IMNS with respect to the fit quality metrics at 7 T and reveals a potential drawback of the IMNS method compared to techniques that aim to fit the MM signals.

The MM peaks identified within the MNS are well in line with resonances of MM signals reported in previous studies^9^.

The correlation analysis indicated 11 and 14 couples of high-correlated MM peaks at 3 T and 7 T, respectively. While p-value adjustment was not applied in this case, a more stringent p-value threshold was chosen to account for multiple testing, which does not exclude Type-I-errors due to potential false positives. However, since the MM ratios are used together as a soft constraint in LCModel, the potential impact of few false positives on the quantification is counterbalanced by the true positives. Moreover, the exclusion of false negative correlations may lead to overfitting, as it results in a higher fit flexibility.

Based on previous studies^8,11,49^ and on the Biological Magnetic Resonance Bank (BMRB)^50^, we assigned the correlations to resonances of the same amino acid (Table 3). It should, however, be noted that MM peaks are rarely investigated and that their assignment is still incomplete^9^.

In this study, the MM peak correlations found at 3 T are mostly different from the correlations found at 7 T. This is not intuitive, as correlations should be based on signals stemming from chemically similar regions, which should be independent of the magnetic field strengths the MMs are subjected to. Some of the differences may be explained by the partial saturation artifact that the MNS exhibit around 0.5 ppm at 7 T. Especially correlations between 0.9 and 1.2 ppm that are assigned to the aliphatic proton resonances of leucine and isoleucine may be affected by this artifact. On the other hand, more correlations were found in the frequency range assigned to α-protons at 7 T. A potential reason for these differences may be a different relaxation behaviour of different parts of the amino acids at different field strengths. However, this needs to be evaluated in future research evaluating MM signal correlations at different echo times and ideally also using different localization sequences.

From previous studies it is known that the baseline estimation has a strong influence on the accuracy of the spectral quantification results^19,51^. Based on observations made in our previous work, and in line with other previous studies ^19, 52, 53^, we chose the baseline spline node distance to be 1 ppm. This provides an appropriate compromise between baseline flexibility and stiffness, as was recommended by Wilson et al.^53^. In the same study^53^ it was found that a too inflexible baseline may lead to a biased quantification. Simicic et al. 2021^19^, on the other hand, observed baseline “distortions” when using a parametrized MM model with low spline node distance values, which decreased when increasing the baseline spline node distances of the parametric MM model to values higher than 0.40 ppm.

We could show that PRaMM exhibits a notably decreased residual and standard deviation in the baseline compared to the other two methods (Figure 4). The baseline of the fit using SCM to incorporate the MM signals exhibits multiple local minima and maxima. It is likely that the smooth and broad shape of MM peaks results in the fact that MM peaks are partly incorporated into the baseline when applying the SCM fit, if the MM intensities deviate from the MM pattern used to generate the SCM basis function. While forcing a flat baseline in SCM might solve this problem, it may introduce bias in the quantification of the metabolites as was demonstrated in a recent study^19^. The PRaMM model, on the other hand, can adapt to altered MM signal intensity patterns, resulting in an increased fit quality, which explains an increased fraction of the spectral signal^54^

Apart from the partial saturation artifact at 7 T described above, the IMNS fits exhibit a smoother baseline than the SCM fits. A bump in the baseline can, however, be observed between 3 ppm and 4 ppm, which may be attributed to a non-optimal MM signal subtraction. Moreover, the residuals show small, structured dips at the frequencies of several main metabolite signals. The culprit for this effect is the non-optimal metabolite-nulling leading to partial subtraction of metabolite signals prior to fitting. Slight signal reduction of a single metabolite peak can partly be compensated using known basis functions, which are fitted to all metabolite peaks of at once. Nevertheless, the quantification results from the IMNS method will be biased by this effect.

Fits based on the PRaMM model exhibit generally an improved quality metric performance compared to the other methods investigated in this study. The average FQNs resulting from fits using PRaMM is lower than FQNs obtained by the other methods in all cases, i.e., the PRaMM model can better explain the data than the other fitting models explored in this study. Although the difference in the FQNs is not always significant, the obtained FQNs are in line with FQNs reported in a parametrized MM model used in rat brain at 9.4T^19^.

While the SNR at both 3 T and 7 T was decreased when using PRaMM compared to SCM, it should be noted that the LCModel SNR is only a rough indicator, as it depends on offsets in the model spectra and baseline^45^. In Figure 4 the SCM fits baseline has a local minimum around 2 ppm. As the spectral maximum is the NAA peak, a local minimum in the baseline at this frequency is bound to increase the LCModel SNR value for these cases. Moreover, in all investigated data sets, the FQN distribution using PRaMM has a smaller interquartile range than the FQNs resulting from the other methods. This indicates that the PRaMM model provides an increased stability of the fit quality compared to the other methods. The increased FQN_nbl_ variability in all three VOIs, compared to FQN_n_, is not surprising. This is because the spectral signal is partially attributed to the fit baseline, which inevitably increases the standard deviation of the residual. This effect is especially pronounced in the case of the SCM fit (see Supporting Information Figure S7), which is likely due to the inflexibility of the MM basis function leading to a higher portion of the MM signal being attributed to the baseline.

Applying the PRaMM model to an unrelated 7 T dataset (RA) resulted in higher performance with respect to all fit quality metrics investigated here, which is a strong indication that the PRaMM model provides a more accurate description of the MM signals in spectral data than SCM.

We found differences in MM peak intensities quantified with PRaMM, both between VOIs and at different field strengths (Figure 6.A,B). Studies investigating the variability of the MM distribution across the brain are rare^23,54–56^. Schaller et al. 2013^54^ found differences in spectral intensity in the frequency ranges 1.5-1.8ppm, 2.3-2.5 ppm, and 3.2-4.2ppm. These differences, among others, are also visible in the data of this study (Figure 6E,F). Nevertheless, no statistically significant differences were found in the MM estimates between the different VOIs, most likely due to the modest number of spectra per VOI. Furthermore, Považan et al. 2018^16^ reported the observation of a positive – although weak - association between the GM fraction and the concentration of various MM peaks in their MRSI study. These findings emphasize the importance to consider the variability of MM signals in different brain regions to achieve accurate MRS quantification.

**Figure 6:**
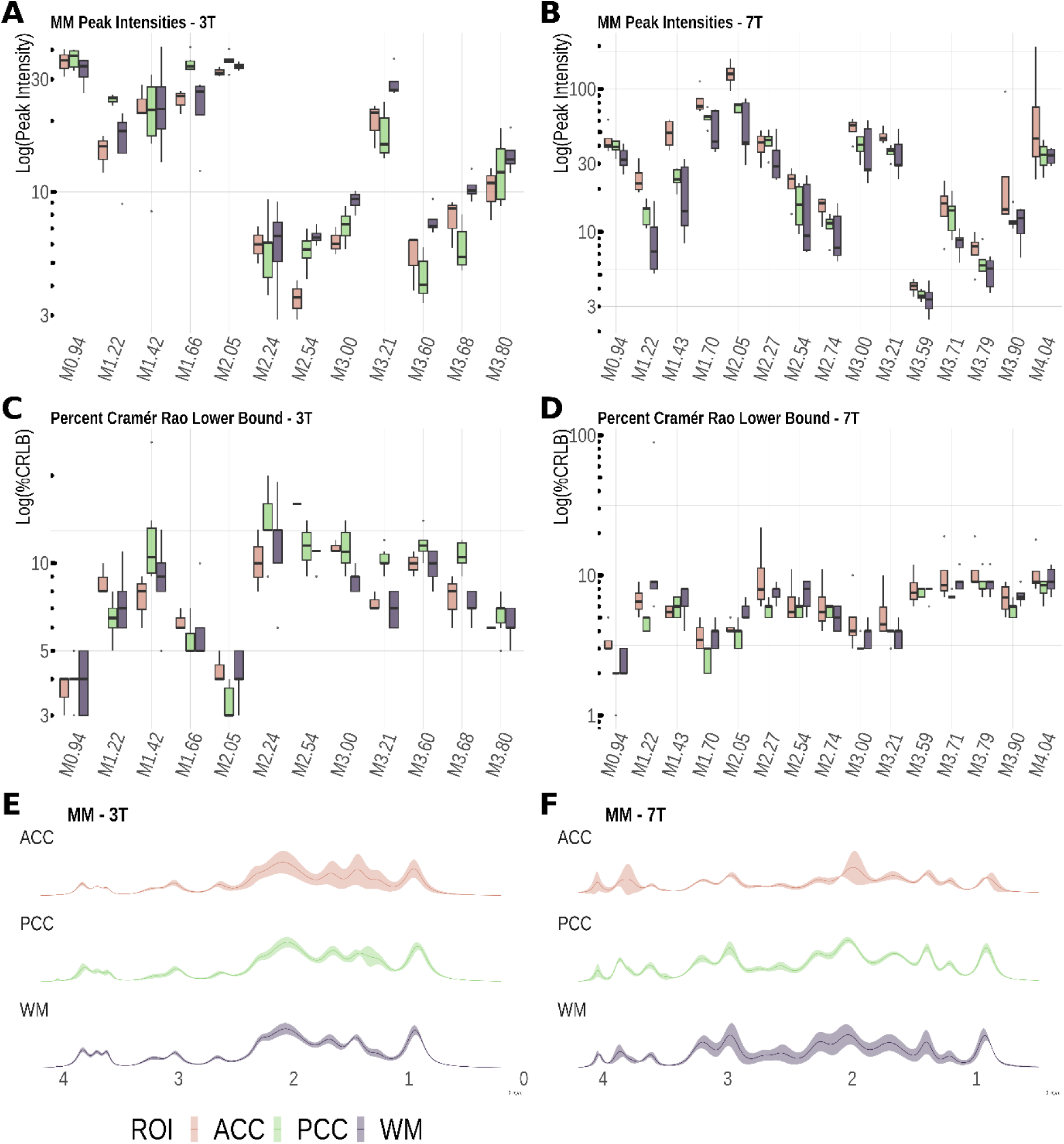
Water scaled macromolecule peak intensities from the fit of the metabolite nulled spectra at 3 T (A) and water scaled and CSF corrected macromolecule peak intensities at 7 T (D) and their differences between the different VOIs, as well as the according relative CRLB%s for both field strengths (B and E, respectively); purple: ACC; green: PCC; and red: WM. E and F display the averaged fits for the respective regions for 3 T and 7 T, respectively, with the standard deviation indicated as shaded areas.

At 3 T and 7 T, similar peak intensities can be observed for the MM signals in the PCC and the WM VOIs, except for M3.21. At 7 T, higher MM levels were, however, observed in the ACC, when compared to 3 T. The most likely explanation for this observation is that due to missing anatomic data the CSF fraction of the ACC VOIs could not be determined at 3 T, and, hence, no CSF correction could be performed. As the ACC is the VOI with the highest CSF fraction among the investigated ones, this effect would be most pronounced here. Another reason may be differing relaxation behaviour between field strength.

Quantification of metabolites revealed more pronounced differences in peak intensities or concentration at 3 T than at 7 T, particularly for major metabolites like NAA and Cr overlapping with major MM peaks at M2.05 and M3.00, respectively. The absolute quantification may have mitigated these differences. Except for metabolites in low concentrations like GABA or GSH, a very a similar pattern is observed at 3 and 7 T: PRaMM seems to quantify slightly higher concentrations in respect to the other models in all clinically important metabolites (Figure 7A,B).

**Figure 7:**
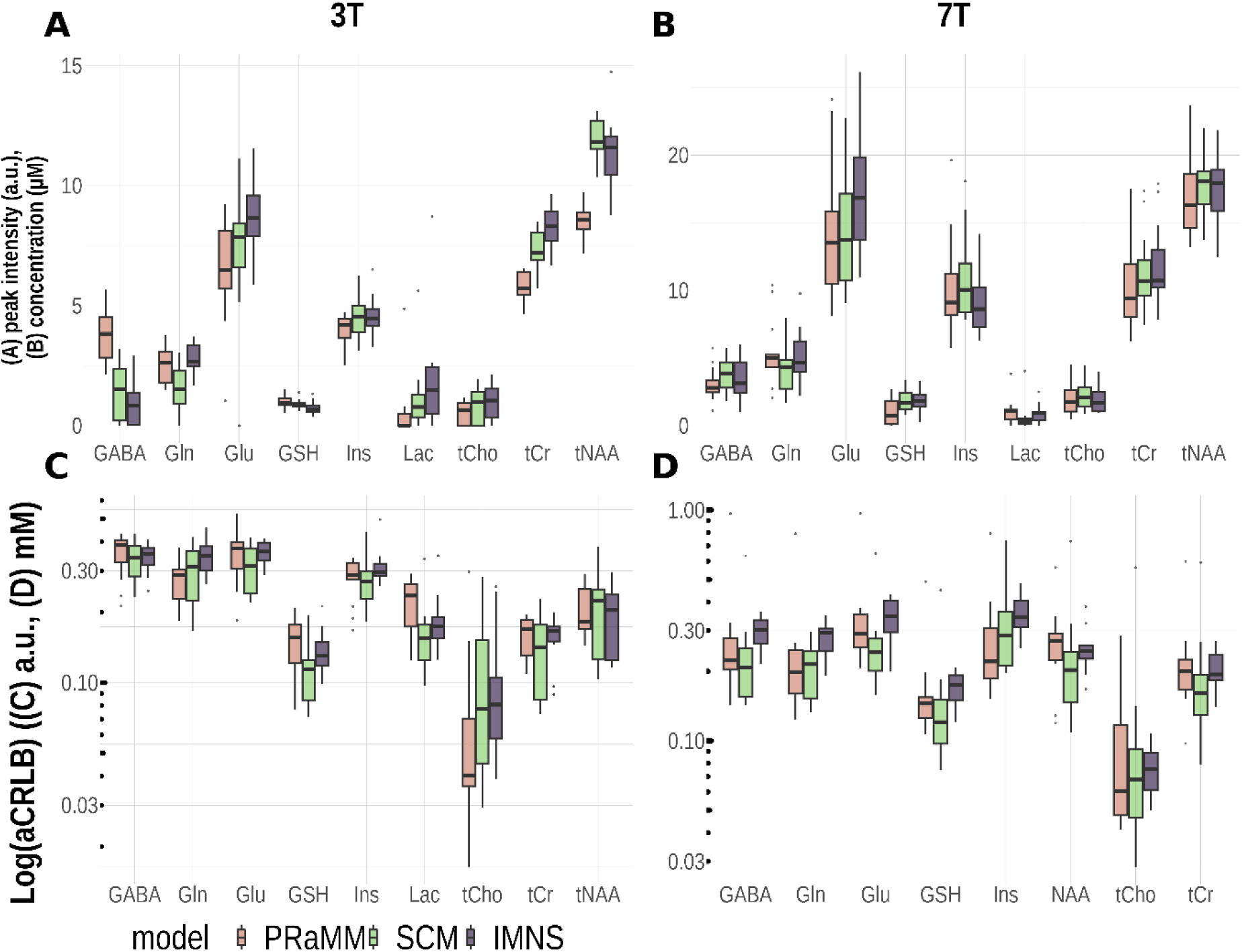
Water scaled peak intensities at 3 T (A) and tissue corrected concentrations at 7 T (B) and their respective relative CRLBs (C,D). It can be seen, that the intensity differences between models are more pronounced at 3T. CRLB differences between models at 3 T are not consistent across metabolites, while at 7 T PRaMM and IMNS have a generally higher CRLB than SCM. For PRaMM this might be attributed to the flat baseline, where it is not possible to account part of the spectral area to the baseline. For IMNS this might partially result from the subtraction of not-fully nulled metabolite peaks together with the MNS

Statistical Testing for differences of the aCRLBs between the different MM models used for quantification showed generally higher values for PRaMM model and lower for SCM (Figure 7C, D). This finding is consistent with that obtained by Heckova et al. 2020^57^,where the parametrized model resulted in higher CRLBs for the main metabolites in multiple VOIs.

Although no statistically significant differences of MM signals in different VOIs were found, it must be assumed that disregarding different MM concentrations in different anatomic regions can introduce bias in metabolite quantification. This has also been pointed out by others^16,54^. Hence, especially for the investigation of rather subtle metabolite concentration changes in neuro-pathologies – or even to establish normal ranges – it is important to find a proper way to treat the MM signals throughout the analysis of spectral data. Whether a parameterized model, such as the proposed PRaMM model, is the way to do this, needs to be investigated in future research.

Analysis of the repeatability and reproducibility on the RA dataset showed no significant effect of the MM model on the metabolite concentrations. While some metabolite concentrations exhibited major differences when different MM models were used for fitting, no model consistently outperforms the other across all metabolites (Figure 8). Whereas the repeatability of the metabolite concentrations appears to be higher when using the SCM, no model outperforms the other when reproducibility.

**Figure 8:**
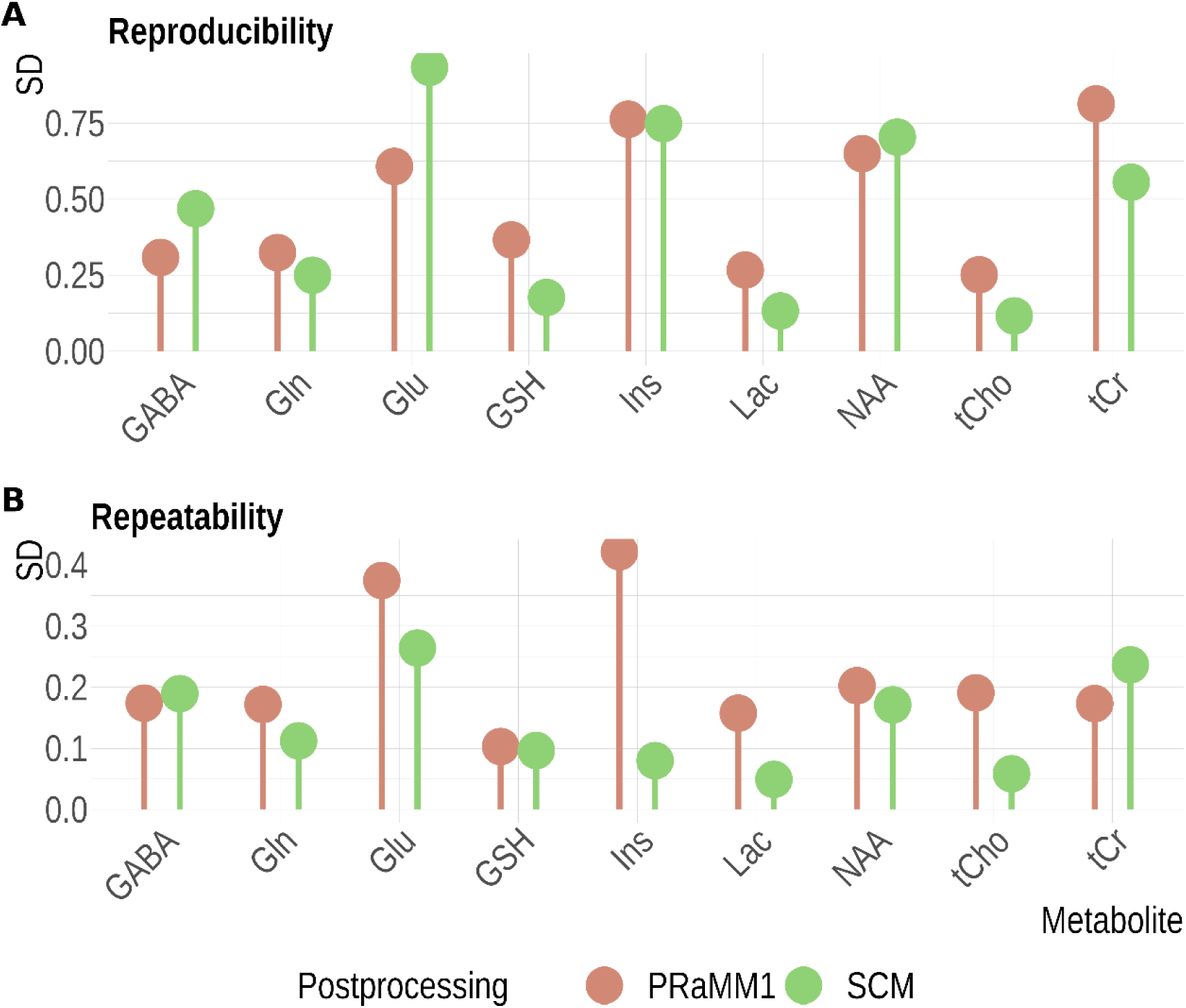
Reproducibility (A) and Repeatability (B) component from REML analysis for all metabolites measured on the RA data set for the use of the PRaMM model (blue) and the SCM (red). SCM overperforms PRaMM with respect to the repeatability, however, with respect to the reproducibility there is no clear trend of one model performing substantially better than the other one.

We found that the CV_%_ values are very good or acceptable for all the clinically relevant metabolites. For almost all metabolites, PRaMM leads to higher CVs_%_ than SCM which is in line with the generally higher aCRLB for PRaMM and the reproducibility and repeatability values on the same dataset.

It is important to notice that SCM, although being more reliable with regard to quantification can only explain a smaller part of the spectrum compared to PRaMM illustrated by the difference of FQNs (Figure 5C and F). It may be thought that part of the spectral area responsible of such uncertainty, that PRaMM account for the metabolite or MM peak area, are accounted by SCM in the fit residual. In absence of a golden standard, it is not possible to distinguish where this part of the spectral area belongs to.

It must be kept in mind that the RA dataset is acquired in the PCC of healthy volunteers and that the SCM makes use of a MM basis function, derived from MN spectra measured in the same region of healthy subjects. As the PRaMM model is based on multiple different VOIs, it is more flexible. It is most likely that the PRaMM outperforms SCM when comparing reproducibility within different VOIs. This needs to be investigated in more detail in the future. Nevertheless, the similar performance of SCM and the PRaMM model regarding reproducibility and coefficient of variation of the metabolite concentrations in the RA data set indicates that the PRaMM model is not overfitting the data, despite adding a lot of flexibility to the fit.

Especially for SCM, the *in vivo* metabolite concentrations in our study are very similar to the results reported by Terpstra et al.^58^, although being obtained in a different study design and acquisition scheme. The CV of metabolite concentrations are, however, may be significantly influenced by variations attributed to normal brain function. Therefore, direct comparisons can only be made if measurements are obtained in the same individual or in larger study cohorts, an essential prerequisite for any application in clinical studies. These differences should be considered in clinical studies, as it is known that measurement uncertainty can distort the results from ANOVA, correlations between variables or other classical statistical methods^59^.

Since MMs do have rather short relaxation time constants, the proper treatment of MM signals throughout the analysis and quantification is especially important when using sequences that allow for short TEs, such as SPECIAL or STEAM. The correlations of MM peak intensities may further be influenced by the choice of localization sequences or changing TE, e.g., in sequences with inherently longer TEs, such as semi-LASER. Furthermore, the effect of the field strength should be investigated in more detail to understand the differing MM peak correlations at

3T and 7T found in this study. In this context, especially the partial saturation that was observed in the MNS in this study should be investigated in future research, as it might have hidden some MM correlations in the 7 T data between two and one ppm, which were found at 3 T. Correlation coefficients are obviously dependent on field strength and TE, why the coefficients of the PRaMM model must be specified according to the localization sequence and the sequence parameters used.

Once a proper treatment of MM signals in the quantification process and measurement uncertainties are consolidated, one of the most important topics of future research will be to establish normal ranges and concentrations of MM signals for healthy tissues and their dependency on the VOI localization. Although we found that differences in metabolite or MM concentrations in different anatomical regions may not be statistically significant, this should be investigated in larger studies with more subjects and multiple VOIs. Finally, the proposed PRaMM model together with a robust multivariate statistical analysis should be applied to establish robust neurochemical biomarkers in different pathologies.

## Conclusion

In this study, the PRaMM model – a novel parameterized MM model using MM peak intensity correlations as soft-constraints – is proposed for improved quantification of proton MRS data. The enhanced fit quality and signal assignment within linear combination modelling has been validated using several fit quality measures. The proposed model enables flexible examination of MM content within different tissues and brain regions. This flexibility could be valuable, especially in situations of potential changes of MM concentrations due to changes over life span or pathological conditions. The risk of overfitting due to the added degrees of freedom was excluded by rigorous investigation of the reproducibility in a data set containing repeated acquisitions within the same healthy subjects. Hence, the proposed model is an important step towards improved characterization of MM in different clinical contexts, starting with the establishment of normal ranges for neurochemical profiles and towards improved diagnosis in several neuropathological conditions.

## Abbreviations in order of appearance

MM: Macromolecule
FS: Full Spectrum/Spectra
MNS: Metabolite Nulled Spectrum/Spectra
PRaMM: Parametrized Ratio Macromolecule Model
FQN: Fit Quality Number
SNR: Signal-to-Noise Ratio
DIR: Double Inversion Recovery
SCM: Single Component Model
IMNS: Individual Metabolite Nulled Subtraction
VOI: Volume of Interest
HLSVD: Hankel Lanczos Single Value Decomposition
SPECIAL: SPin-ECho full Intensity Acquired Localization sequence
WURST: Wideband, Uniform Rate, Smooth Truncation pulse
NAA: N-acetyl aspartate
Glu: glutamate
Gln: glutamine
Asp: aspartate
Cr: creatine
Cho: choline
Ins: myo-Inositol
PCC: posterior cingulate cortex
ACC: anterior cingulate cortex
Ala: alanine
PCho: phosphoryl choline
PCr: phospho-creatine
GABA: γ-aminobutyric acid
GSH: glutathione
Gly: glycine
Lac: lactate
Scyllo: scyllo-inositol
Tau: taurine
Asc: ascorbic acid
BhB: β-hydroxy-butyrate
NAAG: N-acetylaspartylglutamic acid
GPC: glycerylphosphorylcholin
PE: phosphorylethanolamine
Ser: serine

## Acknowledgements

This project (15HLT04 - NeuroMET and 18HLT09 - NeuroMET2) has received funding from the EMPIR program co-financed by the Participating States and from the European Union’s Horizon 2020 research and innovation program. This paper reflects only the author’s view, and EURAMET is not responsible for any use that may be made of the information it contains.

## Conflict of interests

No conflicts of interest to declare.

## Notes

### Competing Interest Statement

The authors have declared no competing interest.

