## Supporting Information for "Macromolecule modelling for improved metabolite quantification using short echo time brain ^1^H MRS at 3 T and 7 T: The PRaMM Model"

### 1. Absolute quantification of the metabolites

For the absolute quantification for each metabolite *m* and each subject *i*, $[H_2O]_{m,i}$, the water signal was considered as the internal standard and its concentrations calculated for each subject as in Equation 1 and Equation 2:

$$[H_2O]_{m,i} = [H_2O]_{pure} \cdot (f_{GM,i} \cdot \alpha_{GM} \cdot R_{H_2O,GM} + f_{WM,i} \cdot \alpha_{WM} \cdot R_{H_2O,WM} + f_{CSF,i} \cdot \alpha_{CSF} \cdot R_{H_2O,CSF})$$

*Equation 1*

$$R_{H_2O,tissue} = \exp\left(-\frac{TE}{T2_{H2O,tissue}}\right) \cdot \exp\left(1 - \exp\left(-\frac{TR}{T1_{H2O,tissue}}\right)\right)$$

*Equation 2*

Where $[H2O]_{pure}$ is the molarity of pure water (55126 mM), $f_{GM,i}$, $f_{WM,i}$, $f_{CSF,i}$ the tissue fractions in the VOI for the volunteer i and $d_{GM}$, $d_{WM}$, $d_{CSF}$ the typical water content fraction in each tissue (0.78, 0.65 and 0.97 respectively)[12]. The resulting value was assigned to the LCModel WCONC parameter.

The output concentrations from LCModel were further corrected for the CSF fraction and the relaxation times as in Equation 3.

$$[m]_{corr,i,v} = \frac{[m]_{i,v}}{\exp\left(\frac{-TE}{T_{2,m}}\right)\left(1 - \exp\left(\frac{-TR}{T_{1,m}}\right)\right)(1 - f_{CSF,i,v})}$$

*Equation 3*

Where $[m]_{i,corr}$ represents the corrected concentration for the metabolite *m* and the volunteer *i* and the VOI *v*; $[m]_{I,v}$ its uncorrected concentration; *TE* the echo time; *TR* repetition time; $T_{1,m}$ and $T_{2,m}$ the longitudinal and transverse relaxation time for the metabolite *m,* respectively; $f_{CSF,i}$ the CSF fraction in the VOI *v* for the volunteer *i.*

To our best knowledge, no $T_2$ values for the MM peaks at 7 T are published yet, for this reason, for 7T and RA the MM water-scaled peak intensities are not CSF and relaxation times corrected.

### 2. Statistical evaluation of the method's precision

The effect of the method on the repeatability and reproducibility of the derived metabolite concentration was estimated by calculating the variance component for each metabolite and method using a restricted maximum likelihood estimation (REML) as described by Riemann et al.[40]. The statistical model was the following:

$$y \sim \mu + S + P + \delta_{session} + \delta_{position} + \epsilon$$

Equation 4

Where y is the observable signal, μ the general mean, S is the fixed subject effect, and P is the fixed post-processing effect. $\delta_{session}$ is the random session effect – nested in subject and preprocessing - and

$\delta_{position}$ describes the random position effect – nested in session – due to little changes in position or scanner calibration during the same-session measurements. Both with mean 0 and variance $\sigma_i^2, i \in \{session, position, \epsilon\}$. The same model is fitted for each metabolite and the variance component is extracted. The significance of the post-processing effects on concentrations was checked via post hoc pairwise comparison – using Tuckey's test - applied to the mixed-effect model. Furthermore, the minimal detectable change (MDC) was calculated using the standard error of measurement (SEM) as in equations (5) and (6). Coefficient Variation (CV) was calculated as well as the ratio of standard deviation and mean within all sessions of each subject. More in-depth about the statistical model can be found in the original paper[20].

$$\mathrm{SEM} = \sqrt{\sigma_{session}^2 + \sigma_{session}^2 + \sigma_{\epsilon}^2}$$

Equation 5

$$MDC = 1.96 \cdot \sqrt{2} \cdot SEM$$

*Equation 6*

*Table S1: Average tissue fractions of the individuals VOIs for the 7 T dataset.*

| | 7T | | |
|---|---|---|---|
| | ACC | PCC | WM |
| GM | 0.48 ± 0.04 | 0.57 ± 0.03 | 0.13 ± 0.03 |
| WM | 0.19 ± 0.03 | 0.29 ± 0.04 | 0.81 ± 0.03 |
| CSF | 0.32 ± 0.06 | 0.13 ± 0.05 | 0.05 ± 0.02 |

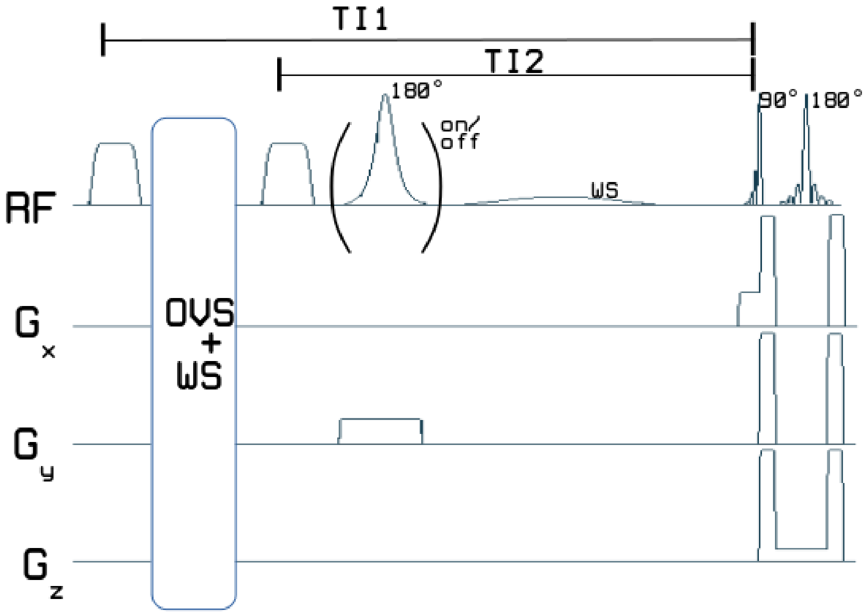


*Figure S1: A schematic representation of the DIR-SPECIAL sequence is provided. To achieve the nulling of metabolites, two WURST pulses were incorporated before the activation pulse, with a customizable distance in milliseconds.*

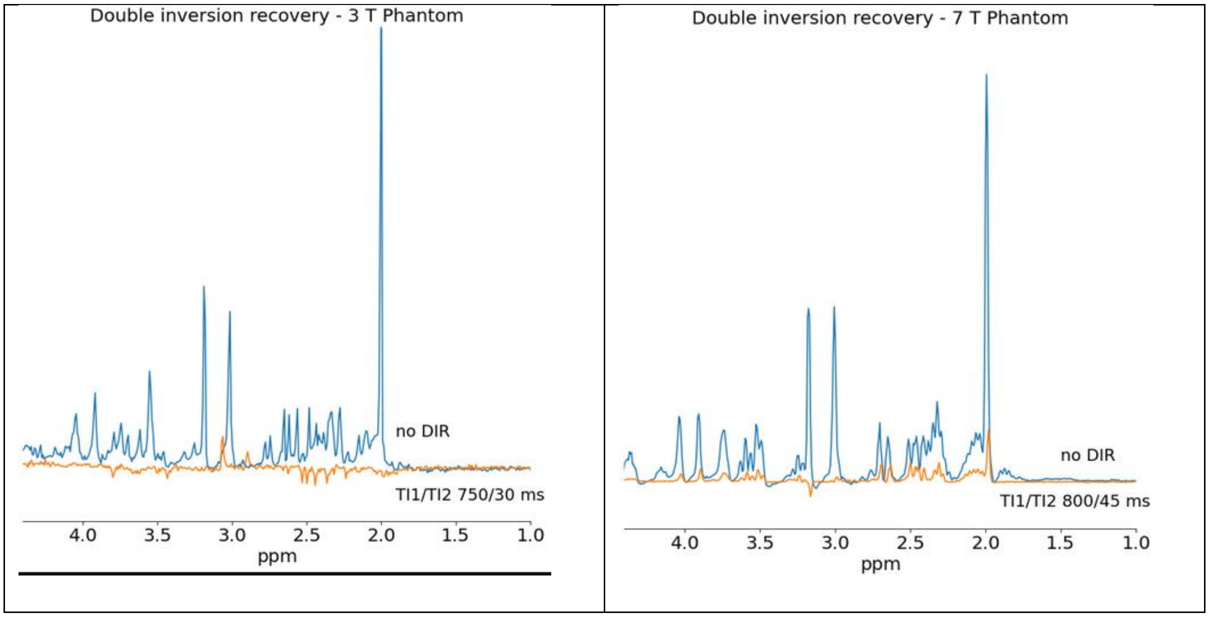


*Figure S2: The full spectrum for the phantom is represented in blue, and the DIR spectrum is depicted in orange for both 3T (A) and 7T (B). It is noticeable that the 7T metabolite-nulled spectrum only contains residual peaks attributed to NAA, Cr, and Glx. In contrast, the 3T MN spectra reveal multiple residual metabolite peaks.*

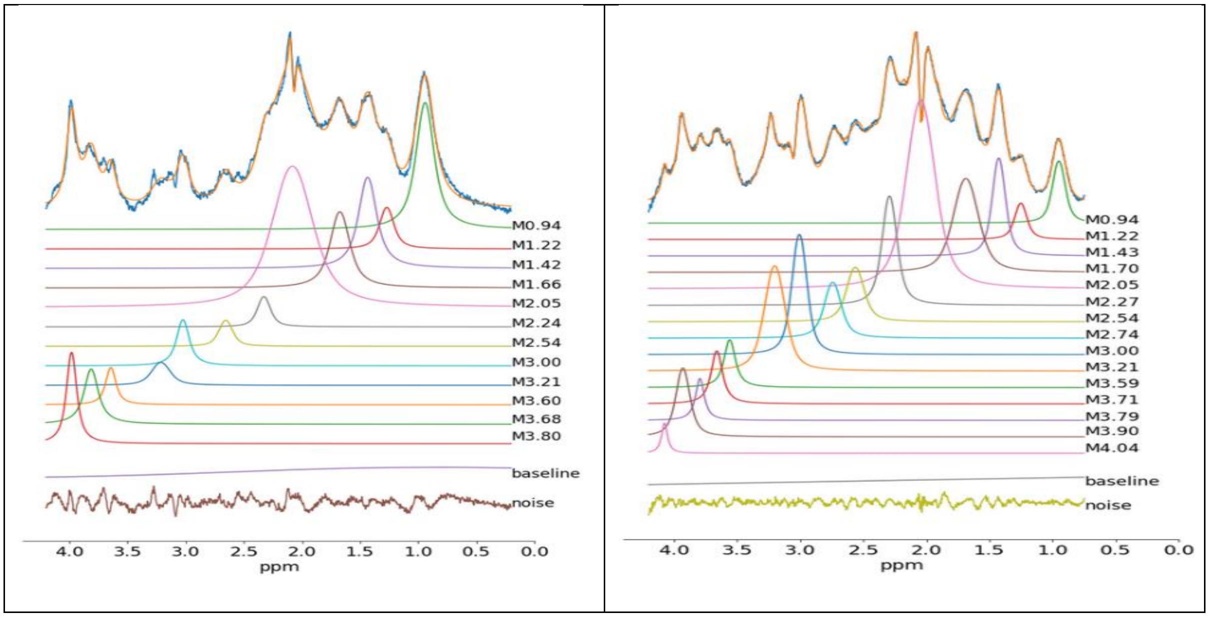


*Figure S3:* MM model and the quantification of the averaged MM spectrum for the 3 T (A) and 7 T

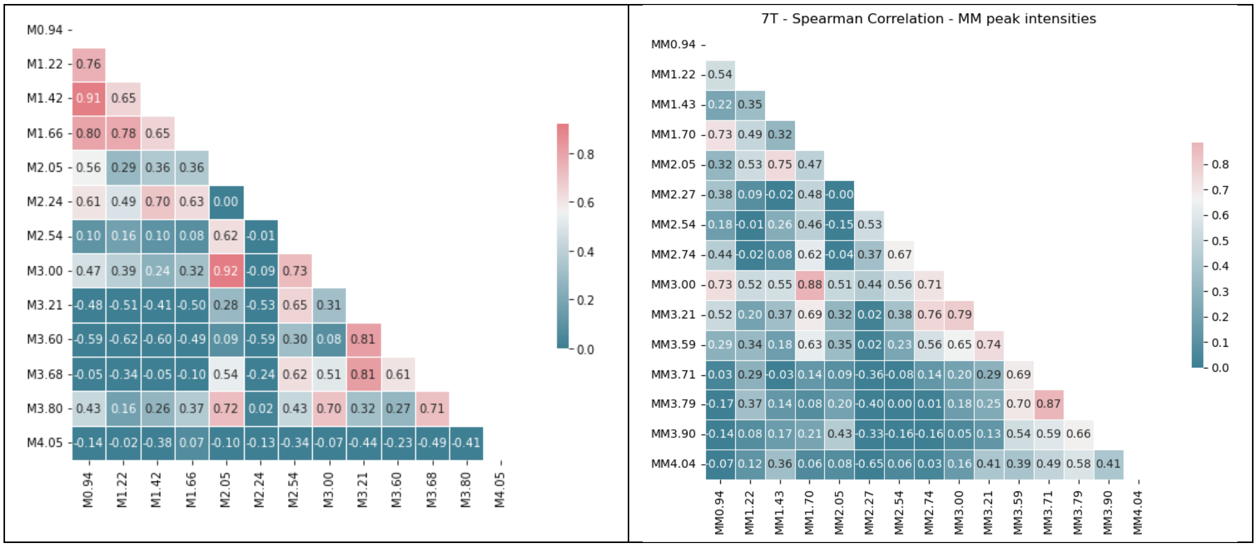


*Figure S4: Correlation matrices for 3T (left) and 7T (right) are presented. The values in the tiles represent Spearman's ρ. Peaks with ρ>0.66 were considered highly correlated. Considering the sample size (N=13), this corresponds to approximately p<0.01.*

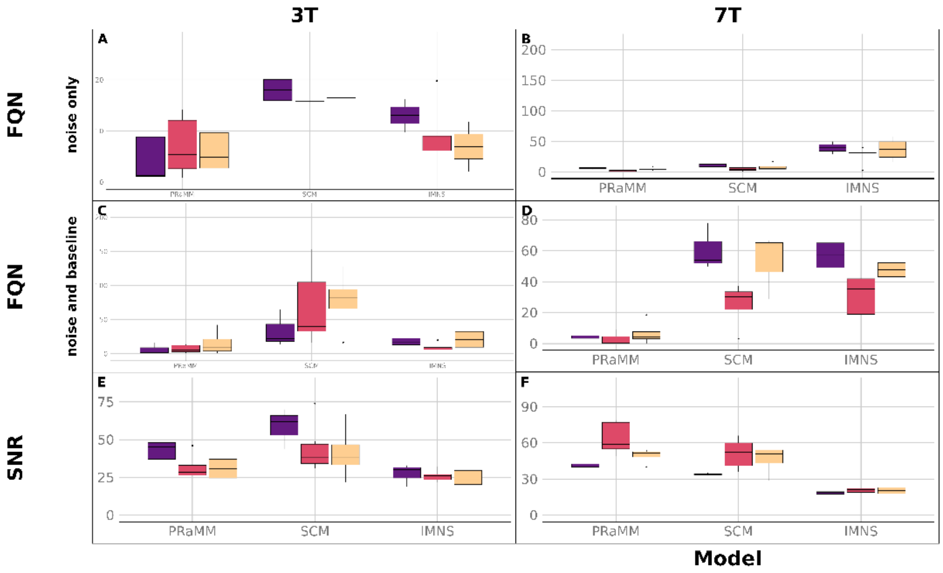


*Figure S5: Fit quality parameters for individual VOIs indicate that, when considering the baseline in the FQN, SCM exhibits a wider range of values, potentially indicative of reduced stability. IMNS demonstrates superior performance at 3T, where optimal MNS subtractions were achievable..*
